# Single-cell proteomics reveals cell-type-specific functional coordination in PBMCs

**DOI:** 10.64898/2026.09.11.751068

**Authors:** Luke Khoury, Andrew Leduc, Saad Khan, Michael Hagemann-Jensen, Jakob Michaëlsson, Jeffrey E. Mold, Nikolai Slavov

**Affiliations:** Department of Bioengineering and Barnett Institute, Northeastern University, Boston, MA 02115, United States; Department of Medicine Huddinge, Karolinska Institutet, Stockholm, Sweden; Department of Cell and Molecular Biology, Karolinska Institutet, Stockholm, Sweden; Parallel Squared Technology Institute, Watertown, MA 02472, United States

## Abstract

The coordination of molecular networks defines cellular functions. This is reflected in molecular covariation within a cell type, which is more subtle than the differences separating cell types and has therefore been difficult to quantify. To achieve the depth, consistency and accuracy required to resolve such covariation, we leveraged single-cell proteomics (plexDIA) and transcriptomics (Smart-seq3xpress) to analyze the proteomes and transcriptomes of thousands of single peripheral blood mononuclear cells (PBMCs). plexDIA quantified more protein-coding gene products per cell with higher data completeness while Smart-seq3xpress quantified more gene products across all cells. The protein measurements revealed cell-type-specific proteome architecture — shaped in part by protein stability, and complex coordination — that was undetectable in our mRNA data. Specifically, protein covariation within a cell type suggests cell-type-specific protein-protein interactions and functional rewiring of biological pathways independent of previously characterized abundance differences. Covariation analysis within cell types resolves a B cell-specific axis of translational states inversely covarying with GDF6 cytokine abundance. Our framework captures cell-type-specific functional coordination representing a distinct information layer accessible via single-cell proteomics.

## Introduction

Peripheral blood mononuclear cells (PBMCs) are frequently analyzed due to their biomedical importance and accessibility. Large single-cell transcriptomic atlases have enabled granular dissection of PBMC subpopulations and differential expression of biological programs across age^1^ and disease^2^ contexts. Protein analysis of individual PBMCs has a long history, from flow cytometry through mass cytometry to DNA-barcoded affinity reagents that enabled the detection of increasing numbers of proteins per cell^3,4^. Still, the number of detected proteins and quantification accuracy remain limited^5^. Transcending these limitations might open a new window towards cellular types and states, especially the functional coordination of proteins^6–8^.

Towards this goal, we used mass spectrometry–based single-cell proteomics that has matured over the past decade^9–17^. These approaches are broadly categorized as label-free or multiplexed using isotopically encoded mass shifts, each balancing trade-offs between proteome depth, accuracy, and throughput^18–21^. Label-free analysis of PBMCs on fast and sensitive MS instruments, such as Orbitrap Astral and timsTOF, can achieve deep and accurate protein quantification, but throughput of sample acquisition (which is set by chromatographic separation time) and depth of proteome coverage trade off sharply^22–24^. Multiplexed single-cell analysis using isobaric mass tags has reached higher sample throughput, but at lower depth and accuracy due to co-isolation interferences^25–27^. To infer quantitative relationships across single cells, we sought to combine quantitative depth, accuracy, and consistency with higher throughput by leveraging the plexDIA framework^28^ and an isotopologous carrier^29^. Such an approach has previously been used to deeply and accurately quantify single-nuclei proteomes^30^ and characterize biological niches within primary tissue^31^.

Here, we focus on extracting a layer of biological information from peripheral blood mononuclear cells (PBMCs) that has remained largely inaccessible to conventional single-cell RNA sequencing: the coordinated covariation of proteins across single cells from the same cell type. This within-cell-type covariation can correspond to protein complexes or functional coordination^32–34^; it is subtler than the variation that distinguishes cell types from one another, and has therefore been difficult to quantify and interpret^35,36^. Yet if resolved, this covariation may reveal important biological structure and protein organization, for example the coordination of physiological functions at the protein level, which has eluded widely used approaches for PBMC analysis.

We applied a 2-plexDIA framework with a 5-cell-equivalent isotopologous carrier using mTRAQ mass tags for single-cell proteomic analysis of human PBMCs, acquired with slice-PASEF^37^ on a timsTOF Ultra platform. Integrating Smart-seq3xpress^38^ transcriptomes from two donor-matched samples enabled quantification of proteins and transcripts across monocytes, NK cells, CD8 T cells, CD4 T cells, and B cells from paired populations. Covariation analysis revealed distinct transcriptome and proteome organization shaped by protein-centric mechanisms, including protein stability and complex coordination, that establish cell-type-specific proteome configurations. Protein covariation analysis between cell types also quantified differential coordination of biological processes independent of mean abundance differences. Finally, within B cells, we resolve an axis of translational-state proteins that inversely tracks GDF6 cytokine abundance, capturing cell-state heterogeneity within a single population. These configurations, and the regulatory information they encode, are largely undetected by single-cell transcriptomic or bulk proteomic methods. Our generalizable covariation framework successfully captures biological information that single-cell proteomics is uniquely powered to quantify.

## Results

### Quantitative characterization and validation of PBMC transcriptomes and proteomes

To characterize the covariation structure underlying cell-type-specific protein and mRNA networks, we collected PBMCs from two healthy donors and profiled paired cell populations by single-cell proteomics and transcriptomics (Fig. 1A). Each dataset was independently projected into a 2D space using independent component analysis (ICA). Both projections clustered cells by types, resolving monocytes, NK cells, CD8 T cells, CD4 T cells, and B cells (Fig. 1B). Similar cell type clustering is observed when using only the 408 proteins quantified in all single cells (Fig. S1A) and without visible clustering of cells by batches and technical factors (Fig. S1C). Cell types were recovered in similar proportions across modalities (Fig. 1C) and between donors (Fig. S1B). Both datasets achieved substantial depth. Smart-seq3xpress libraries detected reads from a median of 4,556 genes per cell across all RNA biotypes (Fig. 1D). Across all batches, plexDIA quantified a median of 14,052 precursor ions per cell, corresponding to 6,690 peptides, and 1,546 proteins (Fig. 1E), substantially exceeding the coverage depth reported previously for primary T cells analyzed by a timsTOF platform^24^.

**Figure 1.**
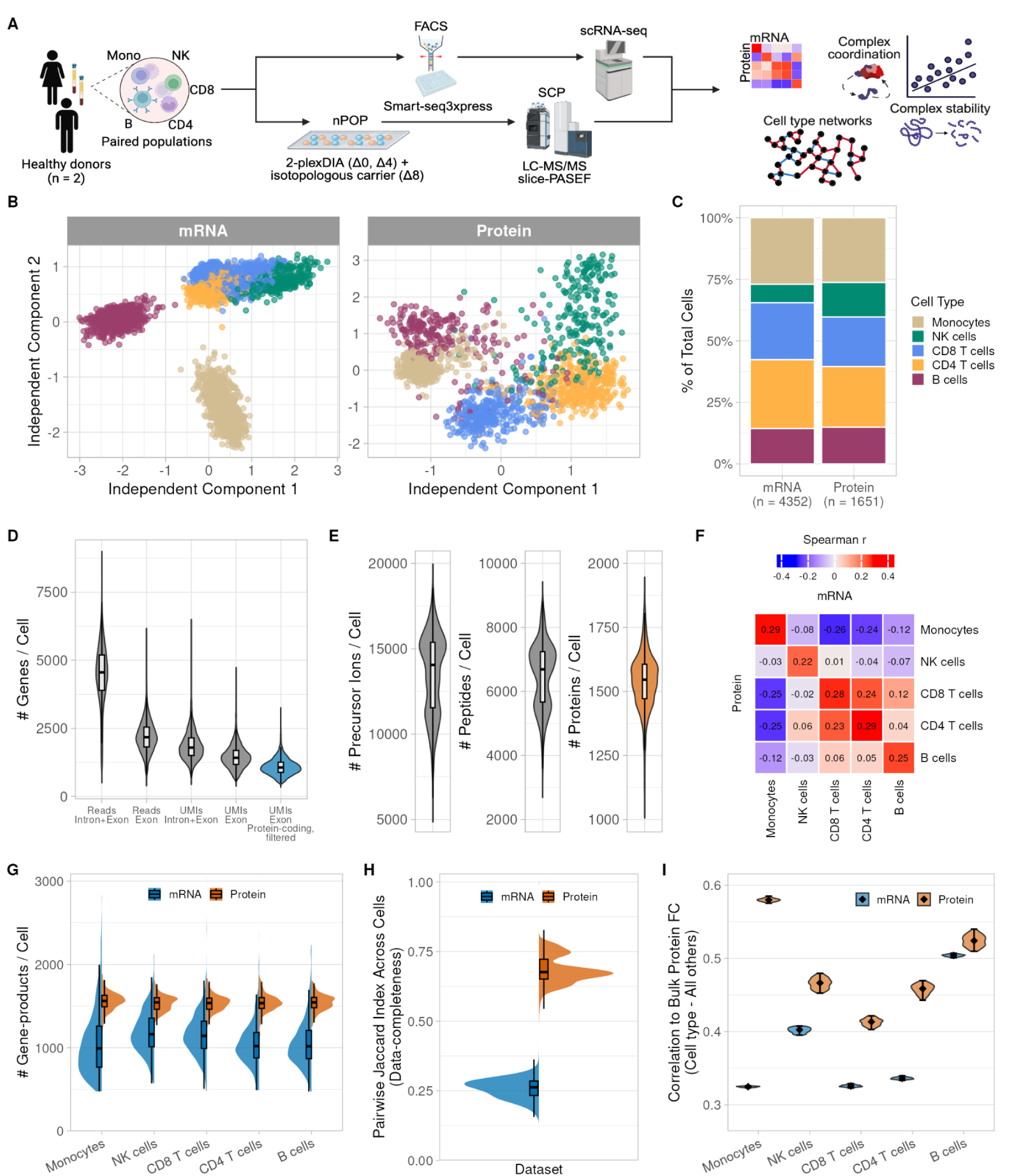
Single-cell proteomics and transcriptomics of PBMCs from donor-matched, paired populations. **A** Data generation pipeline from donor-matched, paired cellular populations analyzed by multiplexed single-cell proteomics with isotopologous carriers and single-cell transcriptomics by Smart-seq3xpress. **B** mRNA and protein single-cell datasets in reduced dimension by independent component analysis, first two components, with cells colored by cell type (3,000 genes, mRNA; 3,732 proteins, imputed). **C** Proportions and numbers of cells per cell type quantified in each modality. **D** Genes from all RNA biotypes quantified by Smart-seq3xpress, filtered for analysis by detectability and abundance for downstream analysis of protein-coding genes specifically. **E** Precursors, peptides, and proteins quantified per cell across all cells. **F** Spearman correlation within and across cell types between relative log_2_ gene-product abundances, computed on cell type means; diagonal entries are within-cell-type mRNA–protein correlations, off-diagonal entries are across-cell-type comparisons. **G** Quantified gene-products per cell across cell types, by modality, restricted to features used for downstream analysis. **H** Data completeness within each modality, estimated as the pairwise Jaccard index across cells over the feature space shared between modalities. **I** Quantitative agreement of 1,254 protein- and mRNA-level gene-product fold-changes, estimated by Pearson correlation of each cell type’s relative log_2_ abundance ratio to the mean of all other cell types, comparing single-cell to bulk protein measurements. Distributions comprise 95% CI of 100 iterations of random 80% subsampling of single cells per fold-change calculation.

The mean relative mRNA and protein levels of cell types defined independently within each modality were compared. The 2,334 gene-products (i.e. transcripts or proteins) detected across all cell types correlated significantly for the same cell type (Spearman r = 0.22–0.29) and much less between cell types (Fig. 1F). A notable exception were the CD8 T cell and CD4 T cell subpopulations, which exhibited only a minor reduction in cross-modal correlation from within-cell-type to across-cell-type (r = 0.28–0.29 to 0.23-0.24) owing to their shared developmental trajectories. mRNA and protein abundances carried markedly different information. Despite similar per-cell feature coverage between modalities (Fig. 1G), the protein data were substantially more complete; pairwise Jaccard indices for 2,651 gene-products detected by both modalities in at least one cell type were significantly higher at the protein than the mRNA level (Fig. 1H), reflecting the high detection consistency conferred by the isotopologous carrier and the plexDIA framework^29^. Throughout, mRNA was quantified as UMI counts of exonic reads from protein-coding genes to compare the pool available for translation, whereas more transcripts were detected per cell when including intronic reads (Fig. S1D).

Cell types were annotated by markers exhibiting differential abundance between resolved cell clusters. Well-established markers showed clear cell-type-specific expression in both the mRNA and protein datasets; flow cytometry-based protein quantitation by index sorting cells used in the mRNA dataset further supported these assignments (Fig. S1E). Having established confident cell-type identities, we further validated our quantitative measurements using multiple approaches including (1) cell types isolated by FACS (Fig. 1I and Fig. S2A), (2) estimating reproducibility of cell-type differences within each modality (Fig. S2B,C), and (3) benchmarking protein abundance across donors (Pearson r > 0.98). Interestingly, while our measurements across all cells detected more distinct transcripts (14,387) than proteins (3,732), the quantified proteins still spanned the transcriptome’s full dynamic range (Fig. S2D).

To assess quantitative accuracy, we spiked a ladder of synthetic yeast peptides into droplets during nPOP^39^ sample preparation for four of five datasets, as previously demonstrated by Huffman et al.^34^. The spike-in concentrations spanned the range of endogenous precursor abundances (Fig. S3A). Regression analysis between measured versus spiked-in concentrations indicated relatively high linearity and precision for the majority of experiments (slopes near or greater than 0.8 and R² > 0.98). To quantify cytokines selectively in the single cells, we added commercially available peptides mapping to relevant cytokines into the isotopologous carrier (JPT Peptide Technologies); an approach analogous to SPIED-DIA^40^ that enables targeted quantification of proteins of interest alongside the global proteome. This yielded consistent detection of intracellular cytokine abundances across single PBMCs in their non-stimulated state, which otherwise would have been difficult to quantify (Fig. S4A). We were then motivated to assess the reliability of protein quantification measurements considering the known challenges of quantifying low abundance proteins, like cytokines. Following Khan et al.^41^, we estimated per-protein quantitative reliability as the median correlation between the mean abundances of randomly split-halves of each protein’s proteotypic peptides across cell types (100 iterations). Both the global proteome and the cytokine subset showed reliability well above a permutation null that shuffled cell-type labels within each peptide, preserving peptide number while removing coherent cross-cell-type signal (Fig. S4B). Cytokines formed a distinctly high-reliability cluster despite their low abundance, comparable to ribosomal protein RPL8, which is far more abundant (Fig. S4C). Reliability also increased monotonically with the number of proteotypic peptides per protein, as expected (Fig. S4D).

Measurement reliability estimation was then extended to the peptide level. Proteotypic peptides from the protein-level analysis were evaluated for how each individual peptide abundance tracks the mean abundance of all other peptides mapping to the same protein across all cell types. This allowed a direct comparison: do peptides from cytokines detected with the spike-in carrier show the same abundances and reliabilities as peptides from the same cytokines detected without it? For the 28 cytokines with peptides in both detection categories, abundances and reliabilities were similar (Fig. S4E), consistent with the isotopologous carrier neither inflating apparent endogenous abundance nor degrading quantitative precision. Peptide reliabilities generally increased with their abundance across all peptides. As at the protein level, cytokine peptides occupied a high-reliability, low-abundance regime (Fig. S4F). Cytokines represent a tightly regulated class of signaling proteins, whose activities may be highly specific and controlled. Regulation may therefore be reflected in their noise (variance) across single cells. Log_2_ protein abundance and its standard deviation were slightly negatively associated (Fig. S4G) across the proteome, with cytokines showing lower than expected variance than the majority of proteins of similar abundance, suggesting their controlled regulation in resting immune cells. Although this analysis does not separate technical from biological variance, the source of that low variability remains unresolved.

### Protein-centric mechanisms underlying biological networks

Cellular functions emerge from networks of interacting molecules, whose coordination is reflected in the correlation of their abundances across cells. Our analyses so far establish accurate and reliable quantification of gene-product abundances across single PBMCs, so abundance variation between cells can be treated as containing biological signal. We used this variation to quantify mRNA and protein covariation within PBMC populations, and asked which mechanisms shape their network structure and cell-type specificity. We find that protein stability and complex coordination organize the proteome in ways largely absent from the transcriptome. At the network level across all gene-products, we observe that protein covariation is largely orthogonal to transcript covariation. Within B cells, protein-protein and mRNA-mRNA correlations were nearly unrelated (r = 0.042; Fig. 2A), consistent with observations in mouse tracheal basal cells^42^ and indicating that transcript co-expression as measured by scRNA-seq poorly predicts protein co-regulation. This decoupling generalized for all cell types; mean concordance across all five cell types was r = 0.052 (SD = 0.011; Fig. S5A) and remained low and stable as networks were restricted to the most reliably measured gene-products.

**Figure 2.**
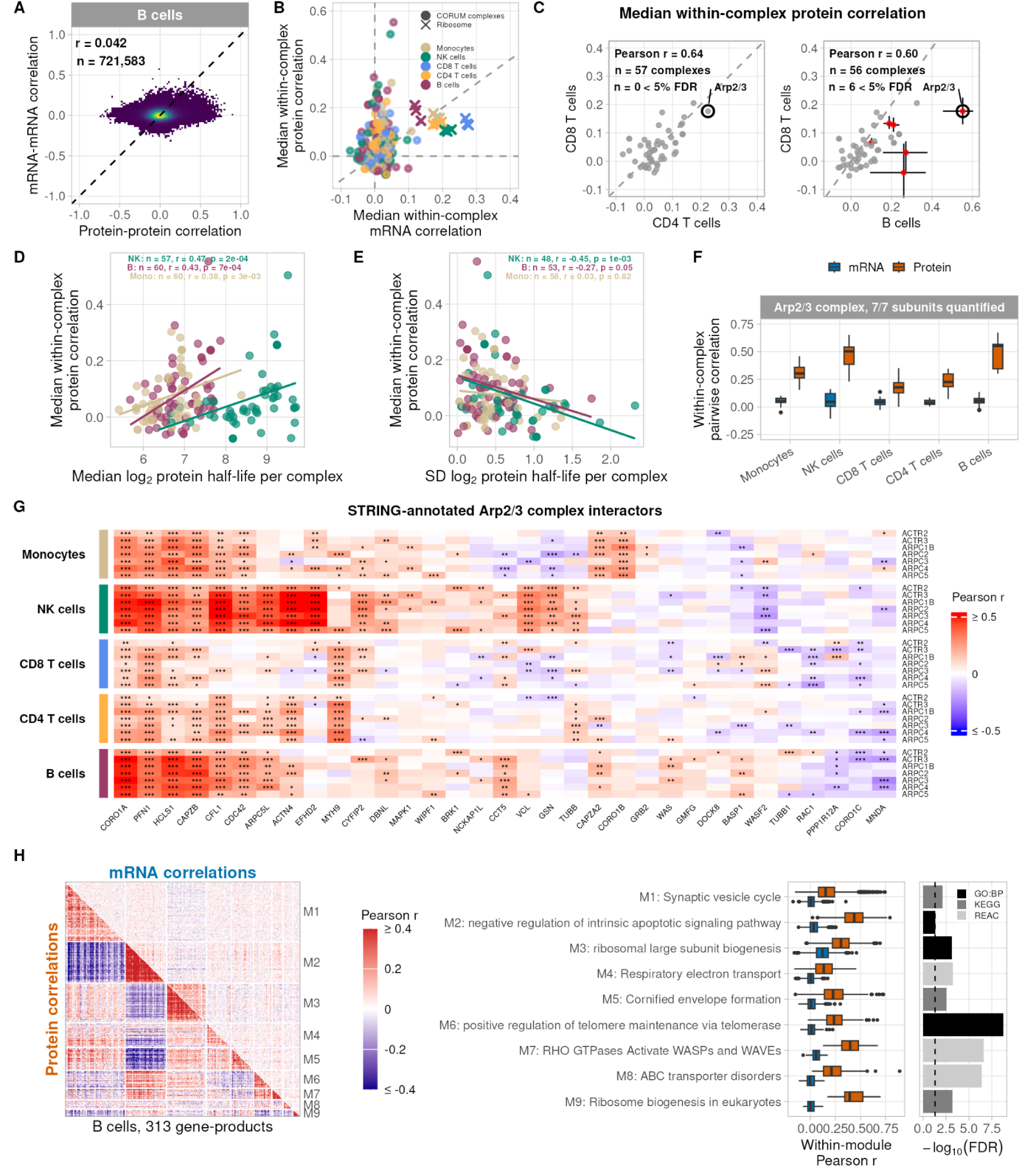
Impact of protein-centric mechanisms on covariation networks. **A** Concordance of mRNA–mRNA and protein–protein correlations within B cells for gene-product pairs shared between modalities (Pearson r = 0.042). **B** Median within-complex pairwise correlations for CORUM-defined protein complexes at the protein- and mRNA-level across all cell types. Ribosomal complexes (80S, 40S, 60S) are annotated. **C** Median within-complex pairwise protein correlations compared between cell types: CD8 T versus CD4 T cells (left) and CD8 T versus B cells (right). Points are medians; error bars are 2.5th–97.5th percentile intervals from 1,000 bootstrap iterations resampling single cells with replacement within each cell type. Between-cell-type differences were formed within each iteration; two-sided bootstrap p-values (twice the smaller tail proportion) were Benjamini–Hochberg corrected across complexes within each comparison. Complexes at <5% FDR are highlighted: six in the CD8 T cell–B cell comparison, none in CD4 T cell–CD8 T cell. **D** Median within-complex pairwise protein correlation versus median log_2_ subunit half-life, from matched-cell-type bulk turnover measurements (REF). **E** Median within-complex pairwise protein correlation versus standard deviation of log_2_ subunit half-lives, per cell type. NK cells show the largest effect size. **F** Pairwise correlations among Arp2/3 complex subunits across cell types, at the protein- and mRNA-level. **G** Correlation structure among Arp2/3 subunits and STRING-annotated interactors across cell types; edges restricted to high-confidence STRING interactions (score ≥ 400). Symbols annotate the percentile rank of each correlation within the distribution of correlations between individual Arp2/3 subunits and 1,543 proteins quantified in all five cell types with at least 100 co-observations. ***, **, and * denote ranks above the 95^th^, 90^th^, or 85^th^ or below the 5^th^, 10^th^, or 15^th^ percentiles, respectively. **H** Protein and mRNA correlation heatmaps for a unified set of highly correlated modules in B cells, defined by hierarchical clustering and annotated by enrichment analysis (GO:BP, Reactome, KEGG).

Next, we sought to identify protein complexes that are coordinated at the protein level in a cell-type-specific manner. Using the CORUM database^43,44^ to define complexes, we computed pairwise correlations among subunits per complex at the mRNA- and protein-level within each cell type. Ribosomal complexes showed strong coordination at both the mRNA- and protein-level across all cell types (Fig. 2B), whereas the majority of other complexes were more coordinated at the protein level. Many complexes are ubiquitously expressed yet serve cell-type-specific functions, assembling subunits under differential stoichiometric constraints. Because such differential assembly should be reflected in how tightly subunit abundances covary across single cells, we compared median within-complex correlations across two cell-type pairs. Between CD8 T cells and CD4 T cells, overall concordance of complex coordination was high (r = 0.64) but no individual complexes differed significantly. Between CD8 T cells and B cells, six complexes were determined to be differentially coordinated at 5% FDR, despite similar overall coordination concordance (r = 0.60). All 6 had higher coordination in B cells (Fig. 2C), indicating cell-type-specific complex coordination.

Complex coordination may be tuned through stability of protein subunits. If stoichiometry is maintained partly through harmonized protein subunit half-life rates, the median within-complex pairwise correlation between subunits should track the half-lives of complex members. Using protein half-life estimates from matched-cell-type bulk turnover measurements^45^, we found that complexes with longer median subunit half-lives are more coordinated across cells (Fig. 2D), suggesting stoichiometry is partially enforced through coordinated degradation. Consistent with this, variability in subunit half-lives within a complex is negatively correlated with coordination, and this relationship is cell-type-specific (Fig. 2E). NK cells show the largest and consistently significant effect, where highly variable subunit degradation rates accompany lower coordination. The association between protein-half life rates and gene-product abundances is protein-centric; within each cell type, protein half-life correlates strongly with protein abundance but negligibly with mRNA abundance (Fig. S5B). These relationships persist within cell types across shared complexes (Fig. S5C, D) and when conditioning on complex size and abundance (Fig. S5E, F).

The Arp2/3 complex illustrates this cell-type-specific, protein-level organization directly. Arp2/3 showed the largest differential coordination between lineages, despite ubiquitous expression and functions in actin-driven cell motility^46^ and membrane organization^47^. Its subunit coordination is cell-type-specific at the protein-level and largely undetectable at the mRNA-level (Fig. 2F), indicating post-transcriptional regulation acting in a cell-type-specific, protein-centric manner. Relating Arp2/3 subunits to their annotated partners through confident STRING interactions^48^ (Fig. 2G), each partner ranked among the strongest 15% of correlations, positive or negative, with at least one subunit in at least one cell type, relative to correlations between that subunit and 1,543 proteins well quantified across all five cell types. Against this background, regulators of Arp2/3 function (CDC42, EFHD2), an alternative complex subunit (ARPC5L), and cytoskeletal filament isoforms (CORO1A, CORO1B, CORO1C) show differential correlation patterns across cell types, indicating that the covariation structure surrounding a single complex is itself remodeled between cell types. These remodeled covariation relationships may reflect known cell-type-specific non-canonical functions of the Arp2/3 complex, such as its role in BCR surface microclustering in B cells^49^ versus its role in cell-cell surface binding for cytotoxic granule delivery in NK cells^50,51^.

Next, we tested whether protein-centric organization beyond annotated complexes can be detected from protein covariation across single cells from the same cell type. To test this hypothesis, we performed correlation module analysis using protein-protein and mRNA-mRNA correlations across B cells. Briefly, gene-product correlations were calculated across B cells for all corresponding mRNAs and proteins. Protein and mRNA correlations, respectively, were hierarchically clustered and modules were identified using the dynamic tree cutting algorithm^52^. Per modality, gene-products were selected for further analysis if they belong to a module with a median pairwise correlation greater than 0.15. The union of all gene-products passing this criteria in either modality were used for recomputing pairwise correlations within each modality, followed by clustering and module identification, yielding a set of highly correlated functional modules defined at the protein- and mRNA-levels. We resolve coherent biological processes within B cells by enrichment across the GO:BP^53^, Reactome^54^, and KEGG^55,56^ databases per module (Fig. 2H). Protein correlation structure revealed strong modular signals that were largely diminished or undetectable at the mRNA-level, providing an interpretable framework for organized proteome function in resting B cells. Clustering the same gene-products by mRNA covariation yielded fewer coherent modules (Fig. S6A). Protein-defined modules also showed greater silhouette widths and median intramodular connectivity than mRNA-defined modules (Fig. S6B), reinforcing prior observations that protein correlations reflect stronger co-regulatory relationships than transcript co-expression across single cells^42,57^. These protein modules are reproducibly observed in each donor’s B cells as shown by the consistent partitioning of cluster members (adjusted Rand index = 0.51, permutation p < 0.001; ARI of zero equates to chance agreement). The strength and specificity of protein correlations are also highly preserved across all modules between donors (Pearson r = 0.43-0.72, permutation p < 0.008) (Fig. S6C, D). Taken together, our single cell protein measurements within a cell type reveal layers of coordinated, pathway-specific biological functions encoded by proteome organization.

### Cell-type-specific protein coordination of biological processes

Our analysis so far indicates that protein covariation captures protein-centric, cell-type-specific information largely unseen at the transcript level. However, it remains unclear whether this organization is captured by differential abundance analysis, which is frequently used to infer functional differences between cell types. To address this question, we first identified clusters of proteins whose abundances covary across single cells within two different populations, monocytes and CD8 T cells. We then compared these within-cell-type covariation patterns to the differential abundance of the same proteins between the cell types.

To define cell-type-specific protein correlation clusters, we used a similar approach done previously for correlation module analysis within a single cell type at the mRNA- and protein-level. Here, we identified shared proteins between monocytes and CD8 T cells and selected for proteins belonging to highly correlated modules within either cell type. For each cell type, we computed protein-protein correlations across this selected set (n = 315 proteins). Each protein therefore has one correlation vector per cell type, describing its covariation with every other protein in that cell type. Correlating the two vectors for a given protein gives a second-order correlation value; a stronger positive value indicates the protein covaries similarly with partners in both cell types, while a negative value indicates it inversely covaries with them. This distribution was broader than an empirical null from randomized data and two-tailed, with each tail representing an extreme of differential co-regulation between the cell types (Fig. 3A).

**Figure 3.**
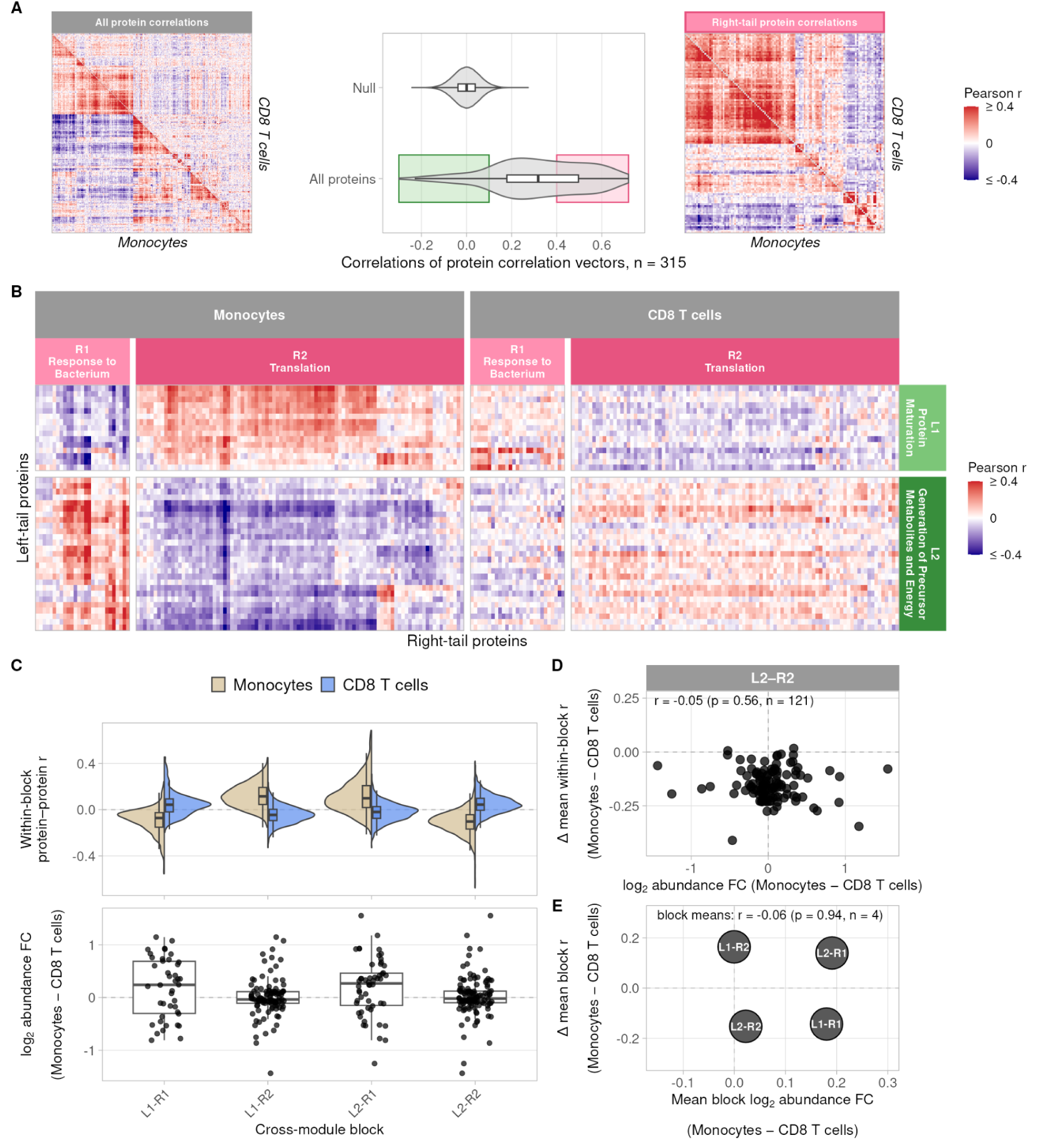
Differential coordination of functional processes between monocytes and CD8 T cells at the protein-level. **A** Protein correlation heatmap (left) for all proteins in analysis (monocytes = lower triangle, CD8 T cells = upper triangle). Distribution of Pearson correlations between the monocyte and CD8 T cell protein–protein correlation vectors (center) for 315 proteins. Right and left tails (r > 0.4; r < 0.1) contain proteins with the most similar and most dissimilar correlation structure between cell types, respectively. Protein correlation heatmap (right) for proteins in the right tail. **B** Proteins from each tail subclustered by hierarchical clustering, annotated by over-representation analysis (GO:BP, <1% FDR), and cross-correlated between corresponding modules within each cell type. **C** Top: distributions of within-module-block protein–protein correlations in each cell type, where module blocks are pairs of correlated protein modules across tails. Bottom: distributions of log_2_ abundance fold change (monocyte − CD8 T) for proteins in each module block. **D** Mean correlation per correlation vector versus log_2_ abundance fold-change (monocyte − CD8 T) for proteins in the L2–R2 module block. **E** Mean values of the metrics in D, correlated across all module blocks.

We resolved these two extremes into functional modules. Taking the proteins from both tails, we computed abundance correlations among them and applied hierarchical clustering with over-representation-based annotation, yielding four annotated modules, two derived from each tail (L1, L2, R1, R2; Fig. 3B). Each block, corresponding to module pairs, showed correlations of opposite signs in the two cell types, indicating that associations between the corresponding biological processes are rewired (Fig. 3C).

However, differences in covariation structure could simply reflect differential protein abundance between cell types. We tested this by comparing mean protein abundances for different module blocks across cell types. Protein abundances differed negligibly between monocytes and CD8 T cells (Fig. 3C) even as the distribution of protein-protein correlations within each block shifted inversely between cell types. Testing this relationship directly, the difference in mean correlation per protein of each block was uncorrelated with its corresponding abundance fold-change (Fig. 3D), which held across all module blocks in the analysis (Fig. 3E). Rewiring of biological networks can thus occur independently of differential abundance. Proteins at similar mean abundance in both cell types can occupy entirely different co-regulatory neighborhoods, which abundance measurements alone cannot resolve. This covariation structure was not observed at the transcript level, suggesting cell-type-specific post-transcriptional regulation driving functional rewiring (Fig. S7A, B). These observations underscore the distinct information layer that single-cell proteomics is uniquely powered to capture.

### Cytokine covariation across B cell translational states

We next tested whether intracellular cytokine abundances reflect cellular states across functional axes within a population. Leveraging the high quantification consistency and reliability of cytokines afforded by the spiked-in isotopologous carrier, we correlated each cytokine against all other proteins within each cell type. GDF6 was significantly anti-correlated with many subunits of the ribosome (Fig. 4A), and this pattern was specific to B cells (Fig. 4B). Only one other cytokine showed a comparable pattern, glucose-6-phosphate isomerase (GPI) in monocytes, which we attribute to its function as an intracellular metabolic enzyme rather than its function as cytokine upon secretion^58^, potentially reflecting a non-cytokine origin for that association.

**Figure 4.**
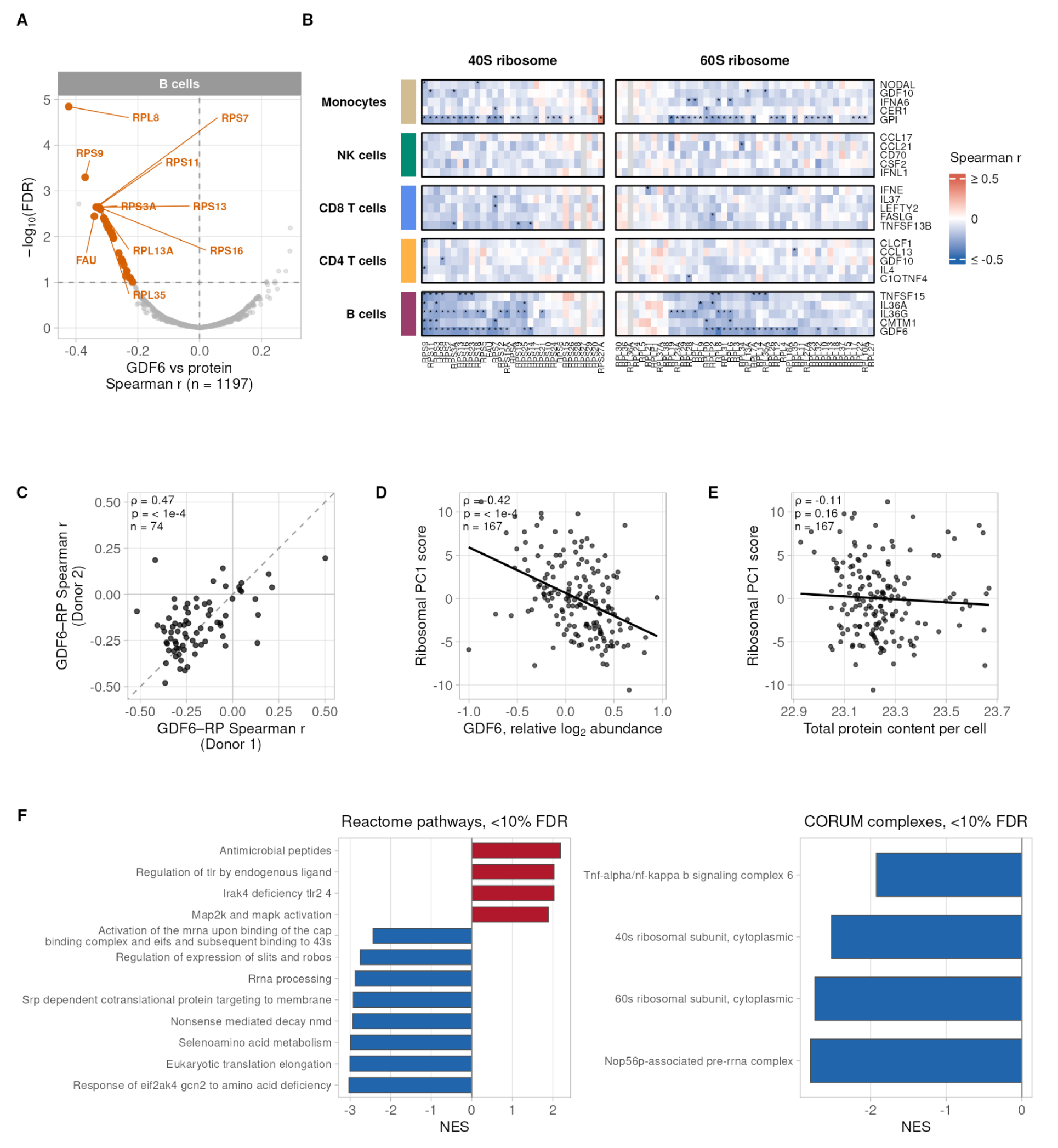
GDF6 anti-correlation with ribosomal proteins across B cells. **A** Volcano plot of correlations between relative log_2_ abundances of GDF6 and each protein across B cells, for proteins with ≥100 co-observations. Correlations computed as Pearson on global per-protein ranks, approximating Spearman. CORUM-defined ribosomal proteins are highlighted; points at ≤10% FDR are orange. **B** Heatmap of correlations between the top-ranked cytokines and 40S and 60S ribosomal proteins across cell types, ranked by number of significant negative correlations and summed negative correlation. Columns clustered hierarchically within B cells. **C** Spearman correlation between per-donor GDF6-ribosomal protein correlation vectors. **D** Ribosomal protein PC1 score per cell, defined as the leading eigenvector-weighted sum of standardized ribosomal protein abundances from PCA restricted to ribosomal proteins, versus GDF6 relative log_2_ abundance across B cells. **E** Ribosomal protein PC1 score versus total protein content per cell across B cells. **F** Protein-set enrichment analysis on the ranked protein list from A, using Reactome (left) and CORUM (right). Terms at ≤10% FDR shown; enrichment direction indicates whether terms are enriched among proteins negatively or positively correlated with GDF6.

The GDF6-ribosome anti-correlation is independent of cell size and reproducible across both donors, shown by the strong positive correlation between per-donor GDF6-ribosomal protein correlation vectors in B cells (Fig. 4C). A single ribosomal-state axis captured the relationship. Per-cell PC1 scores from PCA over ribosomal proteins were negatively correlated with GDF6 abundance across B cells (Fig. 4D). This axis was not a proxy for cell size, as the same PC1 scores showed no association with total protein content per cell (Fig. 4E).

GDF6 marks one end of an axis defined by gradients of translation and immune signaling. To interpret the axis functionally, we performed enrichment analysis on the proteome-wide ranking of GDF6 correlations within B cells. Proteins anti-correlated with GDF6 were enriched for the full translational program, spanning rRNA processing, cap-dependent initiation, elongation, and cotranslational targeting, which were corroborated at the complex level by the cytoplasmic 40S and 60S ribosomal subunits and the pre-rRNA complex (Fig. 4F). The GCN2/eIF2AK4 amino-acid-deficiency response was also enriched at this pole, suggesting the high-translation state is coupled to nutrient stress sensing. In contrast, proteins correlated with GDF6 were enriched for immune signaling, including TLR regulation and MAPK activation (Fig. 4F). GDF6 abundance thus marks one end of an axis of translational states, inversely tracking ribosomal abundance and revealing functional cell-state heterogeneity within resting B cells.

## Discussion

Here, we demonstrate that protein covariation within a cell type encodes distinct regulatory and functional states that single-cell protein analysis can decode across diverse cellular populations. These natural fluctuations in protein abundance may relay the same information that perturbation experiments are designed to reveal^6^, allowing regulatory architectures to be disentangled within a cell type or between conditions by learning how biological networks are structured. Within PBMCs, we find that the underlying structures of transcript and protein networks are largely different, and that protein-centric mechanisms differentially shape how these networks arise.

Our analysis shows at least some of these differences likely reflect protein regulation by degradation (Fig. 2), consistent with previous metabolic pulse analysis of single cells from mouse trachea^42^. However, these differences may also reflect the lower counting statistics and associated signal-to-noise ratio of single-cell RNA sequencing^59,60^, even when using one of the most sensitive transcriptomic methods^38^. Indeed, recent analysis confirms that variation of protein abundance across single cells is dominated by biological signals that far exceed the technical noise^61^.

Protein complex coordination encodes network information specific to the protein-level. We find that subunit stability partially shapes this coordination, consistent with the stoichiometric constraints required for complex function. We further observe that cell-type-specific post-transcriptional regulation can remodel the set of protein-protein interactions surrounding the same complex between cell types. Together, these observations underscore the importance of determining biological context when interpreting the function of a protein or complex. Simply knowing a protein’s abundance may not reveal what the cell is using it for, or how.

Cytokines represent signaling nodes within dynamic pathways and may themselves covary with distinct cellular states. Leveraging isotopologous carriers to selectively quantify these low-abundance proteins, we resolved a B cell-specific axis of translational states covarying with GDF6 abundance. This illustrates how covariation analysis can capture cell-state heterogeneity within a single population that abundance-level analysis alone would not distinguish.

Our covariation analysis yields interpretable biological insight; however, several limitations bound these findings. Although our findings are reproducible across two healthy donors, extending them to a larger and more diverse cohort will be required to establish generality. Our results also reflect network structure in the basal, resting state of PBMCs, which is likely remodeled under perturbation or disease. These measurements therefore provide a healthy resting-state reference against which future studies can compare altered conditions. Our analysis detects associations rather than causal relationships. We mitigated this where possible by using matched-cell-type turnover data to link protein stability to complex coordination, and controlling for donor reproducibility and cell size in the GDF6-translation axis, but these relationships ultimately require controlled perturbation for causal validation. Finally, although the proteome was broadly quantified across cell types, detection sensitivity restricts the depth to which covariation analysis can be applied, and deeper coverage may reveal additional organizational structure.

These limitations motivate clear next steps. As in prior studies that perturb single cells and measure proteome reconfiguration^30^, combining PBMC perturbation with covariation analysis could reveal how initial cell states give rise to different organizational and functional outcomes, helping identify network nodes to target for steering cellular fate. Additionally, single-cell metabolic pulse analysis of PBMCs may clarify translational states within cell types more clearly, as direct synthesis rates may be inferred in these model systems^42,62^. As multiplexing technologies expand, it’s possible to detect covariation structures more precisely by increasing the parallel analysis of cells and proteins simultaneously^63,64^ and improved computational decomposition of multiplexed single-cell spectra^65^. More broadly, protein covariation offers a robust means of interrogating biological systems and discovering functional rewiring between cellular contexts. By capturing a distinct information layer, single-cell proteomics, and in combination with transcriptomics and other modalities, enables a more comprehensive mapping of cellular identity and functional states.

## Methods

### Peripheral blood mononuclear cells

Cryopreserved ampules of human PBMCs from one healthy female (Part # CC-2704, Batch # 21TL228108) and one healthy male (Part # CC-2705, Batch # 21TL132828) were purchased from Lonza. Paired populations of cells were subsequently analyzed by single-cell and bulk analysis methods.

### Antibody staining and flow cytometry sorting

PBMCs were thawed and washed twice in FACSwash (PBS with 2% FCS and 2mM EDTA). The cells were stained with the following antibodies for 30 min at 4°C: CD25 VioBright-FITC (clone 4E3, Miltenyi biotech), CD3 Alexa700 (clone UCHT1), CD4 BUV661 (clone SK3), CD8a BUV395 (clone RPA-T8), CD11c BV605 (clone B-ly6), CD16 APC-Cy7 (clone 3G2), CD19 BUV737 (clone SJ25C1) CD56 PE-CF594 (clone NCAM16.2), CD94 BB700 (clone HP-3D9), FCER1A BV650 (clone AER-37), HLA-DR APC (clone G46-6) (all from BD Biosciences), CCR7 BV421 (clone G043H7), CD123 BV711 (clone 6H6), CD45RA BV786 (clone HI100), CX3CR1 PE (clone 2A9-1) (all from BioLegend), CD14 PE-Cy5 (clone 61D3, Invitrogen), and CD127 PE-Cy7 (clone R34.34, Beckman Coulter). After staining the cells were washed twice in FACSwash and finally washed and resuspended in PBS before sorting. Cells were sorted into 384-well plates (Armadillo High Performance) with index sorting on a FACS Fusion equipped with 5 lasers (BD Biosciences). For cells sorted for RNAseq analysis using Smart-seq3xpess, cells were sorted into 384-well PCR plates containing 0.3ul/well lysis mix overlaid with 3ul/well Vapor-Lock (Qiagen). For proteomic experiments analyzing sorted bulk populations (10,000 cells) of CD8+ T cells (CD3+CD8+), CD4+ T cells (CD3+CD4+), monocytes (CD14+), B cells (CD19+), and NK cells (CD3-CD14-CD19-CD56+) into 96-well PCR plates. After centrifugation, supernatant was removed by pipetting, and the plates were frozen and stored at -80°C until further analysis.

### Smart-seq3xpress library preparation, sequencing, and processing by zUMIs for transcript quantification

Cells sorted by FACS into 384-well plates were processed using the Smart-seq3xpress protocol as described by Hagemann-Jensen et al^38^. Libraries were sequenced paired-end on a NovaSeq 6000 S4 with 10 bp dual indexes, read lengths of 150bp, yielding 450,000 reads/cell. Raw data was processed via zUMIs (version 2.9.7)^66^ for transcript alignment and count quantification. In brief, UMI-containing reads were identified based on the ATTGCGCAATG sequence motif, permitting up to two mismatches. Reads were excluded if the cell barcode contained ≥4 bases or the UMI contained ≥3 bases with Phred scores below 20. The remaining reads were aligned to the human reference genome (hg38) using STAR (v2.7.3a), and read and UMI counts were quantified based on the GENCODE GRCh38 v42 gene annotation.

### Single-cell proteomic sample preparation by nPOP

Single cells were prepared by nPOP^36,39^ for multiplexed proteomic analysis using the 2-plex framework for isotopologous mass tags (mTRAQ; SCIEX). Briefly, synthetic yeast peptides (AYFTAPSSERVEVDSFSGAK and TSIIGTIGPKELYEVDVLK; JPT Peptide Technologies) in LC-MS grade water were dispensed onto fluorocarbon-coated glass slides using the CellenONE system (Cellenion) as previously described by Huffman et al^34^. To ensure quantitative comparability, the peptides were dispensed as an abundance ladder spanning an 8-fold dynamic range, mirroring the precursor ion intensity range of endogenous peptides from single-cells. Droplets were allowed to quickly evaporate before 8 nL droplets of LC-MS grade DMSO were dispensed. Single cells (300 cells/µL 1x PBS) were dispensed into each droplet and, following lysis, were digested overnight using 13 nL of 100 ng/µL Trypsin Gold (Promega V5280) and 20 mM HEPES (pH 8.5), 0.05% DDM (wt/wt) in LC-MS grade water. Peptides from each droplet were then labeled using 20 nL of mTRAQ (Δ0 or Δ4) dissolved in 100% DMSO at 1/60^th^ unit/µL. Paired Δ0- and Δ4-labeled droplets were pooled using an equal percentage of LC-MS grade water and acetonitrile, dispensed into a 384-well plate, dried down via speed vac, and stored at -80°C prior to LC-MS analysis.

### Isotopologous carrier sample preparation

Aliquots of PBMCs from each donor were thawed and washed sufficiently with 1x PBS, resuspended in an appropriate volume, and counted by hemocytometer during the initial steps of nPOP where cells were simultaneously prepared for sorting via CellenONE. Cells were centrifuged, resuspended in LC-MS grade water at a concentration of 2000 cells/µL, transferred to PCR tubes containing 100 µL, and immediately frozen at -80°C. Frozen aliquots were heated at 90°C for 10 minutes per the mPOP protocol^67^ for cell lysis. Lysates were subsequently digested using a final concentration of 20 ng/µL Trypsin Gold, 100 mM TEAB (pH 8.5), and 0.25 units/µL benzonase nuclease (E1014, Millipore Sigma) for 18 hours at 37°C in a thermocycler. Digests were lyophilized and resuspended in LC-MS grade water to a concentration of 2000-cell equivalents/µL. Equal digest volumes from both donors were combined. For datasets incorporating cytokine peptides (SPT-CYT-POOL-hum; JPT Peptide Technologies) in the carrier channel to additionally quantify these proteins of interest in single-cells, approximately 8M copies/cell-equivalent were spiked into the pooled donor digests. To quantify the synthetic yeast peptides including in single-cells prepared by nPOP, these peptides are spiked into the pooled digests at 8M copies/cell-equivalent. Digests were labeled using mTRAQ Δ8 according to manufacturer’s instructions. Specifically, stock concentrations of mTRAQ were added to the digests in volume ratios of 1:2 for label to digest, allowed to react at room temperature for 2 hours with a final concentration of 100 mM TEAB, followed by quenching for 1 hour with 0.1% hydroxylamine. Δ8-labeled samples were lyophilized and resuspended at 2000 cell-equivalents/µL in LC-MS grade water with 0.1% formic acid and 0.015% DDM, then stored at -80°C prior to LC-MS analysis. Carrier samples were diluted to 5 cell-equivalents/µL and used to resuspend single-cell sets in prepared 384-well plates with 1.1 µL.

### Preparation of FACS-isolated populations for bulk proteomic analysis

Plates containing sorted samples were heated at 90°C for cell lysis per the mPOP protocol. Each sample was transferred to a PCR tube using an additional 5 uL of 0.015% DDM in LC-MS grade water, centrifuged, and lyophilized. Samples were reconstituted in 12.3 µL of digestion mix with final concentrations of 16 ng/µL Trypsin Gold, 100 mM TEAB, and 1 unit/µL benzonase nuclease in LC-MS grade water and digested at 37°C for 18 hours. Digests were then centrifuged, lyophilized, and reconstituted in 20 µL of 0.1% formic acid in LC-MS grade water and stored at 4°C until stage tip cleanup. Samples were finally cleaned up to remove excess salts from sorting using a modified EvoTip protocol (Evosep Biosystems). Briefly, prepared tips had samples loaded, centrifuged, washed 3 times with 0.1% formic acid in LC-MS grade water, and collected using 80% LC-MS grade acetonitrile, 0.1% formic acid, and water. Collected samples were lyophilized, stored at -80°C until label-free LC-MS analysis, and reconstituted in 2.5 µL of LC-MS grade water with 0.015% DDM and 0.1% formic acid for acquisition.

### LC-MS/MS

All samples were resuspended in 0.015% DDM, 0.1% formic acid in LC-MS grade water. Bulk samples were injected using 1 µL from glass vials and single-cells were injected using 1 µL from 384-well plates. Different combinations of LC-MS instruments were used over the course of single-cell dataset generation. NanoElute2 (Bruker), UltiMate 3000 (Thermo Scientific), and Vanquish Neo (Thermo Scientific) HPLC systems were utilized. Each chromatographic gradient actively eluted peptides over 30 minutes using 4% to 40% buffer B on 25 cm x 75 µm i.d. IonOpticks Aurora columns with captive spray fittings for timsTOF MS platforms. Bulk analysis utilized the UltiMate 3000 with an active gradient of 60 minutes using 4% to 40% buffer B on the same column type. All LC methods utilized 200 nL/min flow rates. MS analysis was performed using timsTOF platforms, specifically the timsTOF Ultra, timsTOF Ultra 2, and timsUltra AIP, over the course of dataset generation. A modified 1-frame slice-PASEF data-acquisition framework was used for all single-cell analyses, per single-cell methodology described in Sinn et al^37^. Specifically, accumulation and ramp times were set to 200 ms with collision energy set to 20 eV at 1/K_0_ of 60 and 59 eV at 1/K_0_ of 1.60, and collision RF set to 2000 Vpp. From the standard 1-frame method, the MS2 scan range was modified to include 300 to 484 m/z with 1/K0 0.66 to 0.69, 330 to 484 m/z with 1/K0 0.69 to 0.72, 350 to 484 m/z with 1/K0 0.72 to 0.75, and 350 to 484 m/z with 1/K0 0.75 to 0.78. The MS1 scan range was m/z 100 to 1700 and MS2 scan range was m/z 300.2 to 100. The total MS duty cycle was 0.41 seconds. The Athena Ion Processer was turned on for the acquisition of the last single-cell dataset with instrument-specific focus pre-TOF parameters tuned by a Bruker engineer. For bulk analyses and empirical spectral library generation runs, a dia-PASEF method was utilized, as described in Derks et al^30^. Accumulation and ramp times were set to 100 ms with collision energy set to 20 eV at 1/K_0_ of 60 and 59 eV at 1/K_0_ of 1.60, and collision RF set to 2000 Vpp. The MS1 scan range was m/z 100 to 1700 and MS2 scan range was m/z 350 to 1000 with 1/K_0_ from 0.64 to 1.37 and 8 total PASEF frames of 25 Th windows. 4 MS1 scans were used per duty cycle of 1.28 seconds with 1 MS2 scan before every 2 PASEF frames to enable more frequent precursor ion sampling.

### Raw proteomic data processing by DIA-NN

All LC-MS files were searched using DIA-NN^68^ v1.8.2 beta 17. Bulk label-free samples were searched using standard settings against the Swiss-Prot human FASTA (canonical and isoform) with MBR on, mass accuracy set to 15, MS1 accuracy set to 10, scan window set to 6, peak height used for quantification, and the following commands: --peak-translation, -- original-mods, --report-lib-info, --mass-acc-quant 5.0. The second pass results were used for bioinformatic analysis. An mTRAQ-labeled empirical library was generated for searching multiplexed single-cell sets against. First, the same human FASTA had the synthetic yeast peptides used for quantitative accuracy spike-in analysis appended. This FASTA was then used to generate a predicted spectral library using DIA-NN with the fixed mTRAQ Δ0 mass shift of 140.0949630177. Diluted bulk mTRAQ-labeled samples, similarly prepared as described for the isotopologous carrier, were run along with acquisition of each dataset generated. These LC-MS runs in dia-PASEF mode ranged from 100 to 500 cell-equivalents and either contained equal amounts of sample across all three mTRAQ channels or a single mTRAQ channel were searched against this predicted spectral library using the following commands:

--fixed-mod mTRAQ, 140.0949630177, nK,
--channels mTRAQ,0,nK,0:0; mTRAQ,4,nK,4.0070994:4.0070994; mTRAQ,8,nK,8.0141988132:8.0141988132,
--peak-translation
-- original-mods
--report-lib-info

The empirical spectral library containing 7,159 genes, 9,843 protein groups, and 106,881 precursors was used to search each single-cell dataset that was acquired with slice-PASEF. Along with each dataset searched against this same library, diluted bulk mTRAQ-labeled samples run concurrently for QC purposes were also searched with each corresponding batch of single-cell runs to leverage increased coverage through MBR for the second pass results that were used for bioinformatic analysis, as similarly demonstrated by Krull et al^14^. In addition to standard settings, mass accuracy was set to 15, MS1 accuracy set to 10, scan window set to 6, peak height used for quantification, and the following additional commands to those used for empirical library generation:

--mass-acc-quant 5.0
--tims-scan
--tims-stack
--tims-ms1-cycle 3
--ms1-subtract 2
--ms1-base-profile
--tr-ref-profile

### Computational data analysis

#### Single-cell proteomic data processing and integration

Each single-cell proteomic dataset was processed individually then integrated for cell type annotation. Briefly, the DIA-NN report was loaded and cell sorting data from the CellenONE were mapped onto samples using the QuantQC package^39^, with slight modifications for datasets which used an earlier version of the nPOP protocol^36^. Precursors were filtered at ≤1% FDR and protein groups were filtered at ≤5% FDR, and precursors with non-zero MS1 Area were retained. To remove low quality cells and check negative controls, a temporary filter on Channel.Q.Value and Translated.Q.Value at ≤10% FDR was applied. The number of precursors quantified and total MS1 signal was calculated per sample. The median absolute deviation (MAD) was calculated and scaled using a value of 1.4826 for each metric assuming normally distributed data. Cells having less than the median - 1 x MAD for either metric were removed from the analysis. The per-cell median protein CV was calculated for the remaining cells without Channel.Q.Value and Translated.Q.Value applied. This was defined per cell as the median value across all protein CVs calculated each as the CV of normalized peptide abundances mapping to the same protein. The MAD was then computed on this distribution. Cells with median protein CVs above the median + 1 x MAD were removed from the analysis. Protein abundances were calculated using the maxLFQ function from the diann R package^68^ using precursors and MS1 area as input. ComBat^69^ was used to correct for mTRAQ mass tags across cells and, where required for certain datasets, limma^70^ was used to batch correct for LC run order. When required for batch correction, imputation was computed using the kNN algorithm (k=3). Protein abundances were log_2_-transformed. Absolute abundances were normalized by aligning each sample to a reference profile (the row-wise median across all samples) and shifting each sample by the median offset needed to match that reference, correcting for sample-to-sample loading differences while preserving each protein’s absolute abundance scale, as exemplified in Khan et al^41^. Relative abundances were subsequently centered by subtracting each protein’s mean log_2_ abundance across all samples, yielding, for each protein, its deviation from its own mean and thereby removing absolute expression-level differences to isolate sample-to-sample covariation. Peptide abundances were likewise calculated using the same approaches, inputs, batch correction, imputation, and normalization strategies, including the subsequently described dataset integration step.

Protein datasets were integrated by concatenating each protein abundance matrix, yielding parallel absolute (column-normalized) and relative (column-normalized and row-centered) protein matrices, comprising 3,732 proteins and 1,651 cells. Each matrix was normalized, imputed via kNN (k=3), re-normalized, batch-corrected using ComBat with dataset batch as the batch factor, and re-normalized a final time; the original missingness pattern was then restored. Seurat^2^ was used principally for cell type annotation on the integrated relative protein matrix. Missing values were imputed using the row-wise (per-protein) median for this purpose. The top 2000 variable proteins were used for dimensionality reduction, where data were scaled and donor-effects were regressed out. PCA was applied, and the first four principal components were used to construct a nearest-neighbor graph, on which Louvain clustering at a resolution of 0.3 was used for community detection. UMAP was applied using the first four principal components for visualization of cell clusters. An NA-aware Welch’s t-test (one cluster vs. all others, BH-corrected, <5% FDR) was applied to quantify canonical marker protein abundances between cell type clusters using the NA-replaced protein matrix, enabling cell type identification. For visualization of cell types in reduced dimension, independent component analysis was performed on the relative protein abundances, using either a fully kNN imputed matrix or proteins quantified in 100% of single cells (no imputation). All other bioinformatic analyses of protein data utilized the full, non-imputed protein matrix.

### Single-cell transcriptomic dataset processing

UMI counts for exon-only reads from zUMIs processed outputs were loaded for analysis. Protein-coding genes were selected and a Seurat object was created, requiring transcripts quantified in ≥3 cells and cells having ≥200 transcripts quantified. Cells with >500 and <4000 genes and <10% mitochondrial reads were selected. Counts were log normalized using a scale factor of 10,000. To remove sparsely detected and low abundance genes, transcripts were further required to be detected in ≥3% of single-cells have ≥0.05 mean log-normalized expression, retaining 7,098 genes (49.2%) and 4,520 cells. Following quality control, single-cell expression data was used for dimensionality reduction and clustering for cell type annotation. Briefly, the top 3,000 variable genes were selected using variance-stabilizing transformation. Donor effects were scaled and centered, during which donor-effects were regressed out. PCA was applied, and the first 50 principal components were used to construct a nearest-neighbor graph, on which Louvain clustering at a resolution of 0.9 was used for community detection. UMAP was applied using the first 50 principal components for visualization of cell clusters. Differential expression testing using Welch’s t-test (one cluster vs. all others, BH-corrected, <5% FDR) was applied to quantify canonical marker gene abundances between cell type clusters, enabling cell type identification. For clusters in which subpopulations could be resolved as subclusters, the main population was annotated for each subcluster, and the differential expression analysis was repeated using the main population clusters for final visualization. During FACS-isolation of single cells during Smart-seq3xpress sample preparation, a small minority of samples did not have FACS measurements due to data collection issues. The remaining majority of cells that had paired FACS-based protein abundances quantified were used to validate the cell type assignments. Fluorescence intensities for proteins were transformed using arcsinh and a cofactor of 150 to compress the bright tails and stabilize variance. Each marker protein was z-scored across cells and detection of positive cells per marker was set as those with intensities ≥ median + 1 standard deviation. Dendritic cells were removed for all subsequent analyses, since they were not identified in the single-cell protein dataset. To quantify detection sensitivity from the Smart-seq3xpress dataset, the number of genes per cell from the set of selected cells from zUMIs processed count matrices were calculated, including read counts or UMI counts for either intron and exon-containing reads or exon-only containing reads. For visualization of cell types in reduced dimension, independent component analysis was performed on the scaled expression data.

### Bulk proteomic data processing

The DIA-NN report for bulk protein data was loaded and precursors were filtered at ≤1% FDR and protein groups were filtered at ≤5% FDR, and precursors with non-zero MS1 Area and MS2-level fragment ion intensity were retained. Precursors were then required to be detected in ≥2 replicates per cell type and in ≥2 cell types across the dataset. Protein abundances were calculated using maxLFQ and normalized as previously described. Samples were visualized in reduced dimension by independent component analysis on the set of 346 proteins quantified in all single-cell proteomes that were intersected with bulk proteomes from FACS-isolated subpopulations.

### Assessing quantitative accuracy using synthetic yeast peptide spike-ins

Synthetic yeast peptides were spiked-in at the beginning of each nPOP protocol in four of five datasets using an abundance ladder as previously described to assess the accuracy and precision of quantification per dataset. Precursors mapping to either the synthetic yeast peptides or endogenous human peptides were annotated. Precursors were filtered at ≤1% FDR and protein groups were filtered at ≤5% FDR, and precursors with non-zero MS1 Area were retained for analysis. Further, spike-in precursors were selected if quantified in >70% of samples per abundance level. Within each abundance level, spike-in precursors with quantities below the median – 1 x MAD were excluded to remove low-quality signal attributable to variable dispensing volumes of 300 pL using the CellenONE. CVs were calculated per precursor per spike-in level after normalizing each sample to its column mean, then averaged across precursors within each level to obtain a mean CV per level. To quantify measurement accuracy, each precursor’s abundance was normalized to its own median abundance at the lowest spike level as a reference, then summarized as the median normalized abundance per peptide sequence (collapsing across charge states) at each spike level. A robust linear regression (rlm) was fit between log2-transformed expected spike-in levels and log2-transformed median peptide measured abundance per level, and the fitted slope and coefficient of determination (R²) estimated as the squared Pearson correlation were used to evaluate quantitative accuracy and dynamic range across the dilution series. Precursor-level abundances of retained spike-in peptides were compared against those of endogenous human precursors across samples to contextualize spike-in signal relative to single-cell abundance range. Robust regression of measured versus known abundances yielded slopes at or above ∼0.8 and high goodness-of-fit (R² > ∼0.98) for three of these datasets; the fourth used an earlier nPOP version^36^ later optimized to reduce surface losses^39^, which may have affected recovery of these spike-in peptides due to multiple drying steps.

### Protein quantification reliability estimation

Protein quantification reliability was assessed using methodologies described by Khan et al^41^. The consistency of protein abundance variation across cell types can be estimated using the mean abundance of randomly split sets of peptide relative abundances mapping to the same protein. This analysis was restricted to proteotypic peptides, requiring that peptides were quantified in ≥10 cells within each cell type, and proteins must have ≥2 peptides passing these thresholds to be included in the analysis. For each protein meeting these criteria, a randomized, non-overlapping split-half set of peptides are obtained, and the mean relative abundance is calculated for each cell type. The independently calculated means are correlated using the Pearson method across cell types to determine the split-half correlation. This procedure is repeated 100 times, and the median split-half correlation is taken as the protein’s reliability estimate. To establish an empirical noise floor, a strict permutation null was generated by shuffling cell-type labels independently within each peptide, preserving each protein’s peptide count and margin value distribution while destroying real cross-cell-type signal, and recomputing the split-half correlations using the median correlation across 100 iterations as described. Peptide-level reliability estimates were similarly calculated using a modified framework. For each peptide a leave-one-out (LOO) statistic was calculated, defined as the per-cell-type mean profile correlated against the mean profile of all remaining peptides mapping to the same protein across cell types.

### Concordance analysis of mRNA and protein covariation networks

Relative log_2_ mRNA and protein abundances were used as input into the concordance analysis pipeline. Pearson correlation with pairwise complete observations was first used to calculate mRNA-mRNA correlations and protein-protein correlations for the shared set of gene-products within each cell type individually, requiring gene-products to have non-zero variance in both modalities. The set of protein-protein correlations was then filtered such that ≥100 co-observations were used to calculate the concordance. This filtered set of gene-product pairs was intersected with the mRNA-mRNA correlations for concordance between these covariation networks using the Pearson method. This is repeated for each cell type and the mean value and standard deviation is calculated, followed by the same process across cell types using a filtered set of gene-products having increasingly higher positive protein-level quantification reliability estimates using steps of 0.1. For each step, the mean number and standard deviation of correlation pairs within the concordance estimates are calculated across cell types.

### Cell-type-specific protein complex and stability coordination analysis

Protein turnover data from Mathieson et al^45^ was used for the matching cell types (Monocytes, NK cells, and B cells). Protein half-life measurements were retained where R² was ≥0.80 and the per-replicate quality flag was "good" across all replicate columns. Replicate half-lives were then averaged within each cell type. The CORUM database (version 5.2)^44^ was used to define protein complexes with analysis performed on complexes with ≥3 protein subunits. For each protein complex within each cell type, protein subunits were required to be quantified in ≥70% of cells and complexes were only analyzed further if it had ≥70% of its annotated subunits quantified in both the protein and mRNA datasets. The median pairwise correlation between all complex subunits per complex was calculated for each cell type at the protein and mRNA level. Cell-type-resolved complex coordination was determined for CD4 T cells, CD8 T cells, and B cells by calculating the median within-complex pairwise correlation for each complex using 1,000 iterations of a non-parametric bootstrap of single-cells with replacement per cell type and taking the 2.5^th^ to 97.5^th^ percentile intervals of these bootstrap distributions as error bars. To test whether a complex exhibited cell-type-specific coordination, the difference in median correlation between the two cell types was formed within each bootstrap iteration, yielding a bootstrap distribution of the between-cell-type difference per complex. A two-sided bootstrap p-value was computed as twice the smaller tail proportion of this difference distribution and p-values were BH-corrected across complexes within each cell-type-pair comparison. Complexes with corrected <5% FDR were considered to show statistically resolvable cell-type-specific within-complex coordination and are highlighted. Among complexes across NK, B, and monocyte cell types, the median, and separately the standard deviation, of subunit log_2_ half-life per complex was related to the median within-complex protein correlation by Spearman correlation. To distinguish a half-life effect from confounding by complex size and abundance, partial Spearman correlations were computed controlling for the number of detected subunits and for median subunit log₂ abundance (ppcor package^71^). To quantify cell-type-specific covariation of the Arp2/3 complex with its known protein-interaction partners, the STRING database^48^ (version 12.0, human, physical subnetwork) was queried for protein-protein interactions with combined-score threshold of ≥400, restricting edges to those supported by experimental or curated-database evidence. Each subunit of the Arp2/3 complex was correlated (Pearson) using pairwise complete observations with all interaction partners within each cell type, reporting only those with ≥100 co-observations within each cell type. To express the magnitudes of these correlations on a scale interpretable across subunits and cell types, a background set of proteins was determined independently from STRING-based annotations. The set of proteins quantified in at least 40% of cells within each cell cell type were taken, excluding Arp2/3 subunits, and were subsequently filtered to keep only proteins that had at least 100 co-observations per correlation with each Arp2/3 subunit per cell type. The final set of 1,543 proteins was taken as the intersection across all cell types. Correlations between each subunit and these 1,543 proteins were then recomputed within each cell type, and correlations for each STRING-annotated protein were ranked among the resulting distribution of correlations per Arp2/3 subunit, per cell type. Ranked correlation percentiles were denoted either either ***, **, or * for correlations above the 95^th^, 90^th^, 85^th^, or below the 5^th^, 10^th^, or 15^th^ percentiles, respectively.

### Correlation-module analysis

Within B cells, gene-products detected in at least 60% of cells in the protein dataset and simultaneously present in the mRNA dataset, with non-zero variance in both modalities, were retained. Pearson correlation matrices were computed for each modality, converted to distances (1−r), and hierarchically clustered (Ward’s D2^72,73^). Modules were defined by adaptive branch cutting^52^ (dynamicTreeCut, hybrid method, deepSplit 1.5, minimum module size 10) and retained only if their median within-module correlation was ≥ 0.15. The union of protein-derived and mRNA-derived module genes was re-clustered to define the final module set. Functional enrichment of each module was assessed with g:Profiler^74^ (GO:BP, KEGG, Reactome) against the analyzed gene set as a custom background with FDR correction ≤10% as the upper limit. Terms were selected as having the lowest FDR between three tiers (less than 1%, between 1% and 5%, and between 5% and 10% FDR) with the largest effect size (intersection size/term size), and ties broken by choosing the lowest FDR value within the best tier. Module quality was quantified per modality by silhouette width and by intra- versus inter-module connectivity (pairwise correlations). Cross-validation of protein modules identified from the same space of gene-products from the primary analysis were identified independently in Donor 1 (n=122) and Donor 2 (n=124) B cells using the same procedure (Pearson correlation, Ward’s D2 clustering, dynamic tree cutting) using 309 of 313 proteins above the detection threshold in both donors. Partition agreement was calculated using the adjusted Rand index, with significance determined by 1,000 permutations of module labels. Comparison of correlation module magnitudes and structure were assessed between donors using the protein modules defined in Fig. 2H so that protein clusters were identical. Protein pairs with fewer than 100 co-observations in either donor were excluded. Correlation structure preservation was computed by correlating the pairwise complete set of vectorized protein-protein correlations between donors within each module, with significance calculated from 1,000 permutations of protein labels within each module.

### Differential protein covariation between cell types

Analysis was restricted to proteins quantified in every Monocyte and CD8 T cell across both populations. Pearson correlation among these proteins within each cell type were calculated, and hierarchically clustered (Ward’s D2 on 1 − r) with adaptive branch cutting (dynamicTreeCut, hybrid method, deepSplit 3, minimum cluster size 5), retaining only proteins in modules whose median within-module correlation was ≥0.2. The union of proteins from modules passing this threshold from both cell types were then selected for correlation vector analysis and empirical null generation. Briefly, within each cell type, all pairwise Pearson correlations among these proteins were computed. For a given protein, its vector of correlations to all other proteins in one cell type was correlated (Pearson) against the corresponding vector in the other cell type. For each protein, we correlated its corresponding correlation vectors, and the resulting correlation served as its rewiring score with a low value indicating that a protein’s covariation partners differ between cell types (differential covariation), and a high value indicating similar covariation. The empirical null was generated by independently permuting the values within each cell-type’s matrix and recomputing rewiring scores over 1,000 permutations. From the distribution of correlations of correlation vectors, proteins were partitioned into two tails defined as the differentially-covarying left tail (≤ 0.1), and a similarly covarying right tail, (> 0.4). Cross-set correlation matrices between the left tail and right tail proteins were then computed within each cell type. Using the monocyte cross-set as the reference, the left and right tail proteins were each split into two clusters by Ward’s-D2 hierarchical clustering (k=2). These monocyte-derived cluster assignments (L1/L2 and R1/R2) were carried over to the CD8 T cell matrix so that both cell types were organized using the same correlation matrix ordering for comparability. Each of the four tail-derived clusters was tested for GO:BP over-representation (MSigDB^75,76^ C5 GO:BP gene sets) using an over-representation test (fgsea package^77^) against the universe background defined as the set of proteins input to the correlation vector analysis, retaining terms at <1% FDR with gene-set sizes between 3 and 300. Cross-module blocks defined as those correlation submatrices between tail-cluster pairs were then used to define blocks for further analysis. For each block, the mean within-block correlation was computed per cell type and its between-cell-type difference (monocyte – CD8 T cell) was calculated, representing the magnitude of differential covariation between cell types. Protein abundance fold change between the two cell types was computed as the difference in per-cell-type mean abundance. Pearson correlation was then calculated between within-block differential covariation magnitude and protein abundance fold change, at the per-protein level within a block and across block-level means. This analysis was reapplied at the mRNA-level using the same set of gene-products and monocyte-derived cluster assignments. The corresponding within-cell-type and cross-set correlation heatmaps were generated for comparison against the protein-level structure.

### Cytokine covariation analysis

Analysis was performed in the space of cells measured from the three datasets where the cytokine panel of peptides was included in the isotopologous carrier. Within each cell type, cytokines quantified in ≥100 cells were correlated pairwise with all non-cytokine protein quantified in ≥40% of cells using a fast approximation of Spearman’s method where values were converted to global per-protein ranks and correlated (Pearson) using pairwise-complete observations, retaining correlations with ≥100 co-observations for further analysis. Two-sided p-values were derived from the correlation and the co-observation count and BH-corrected for multiple testing separately within each cytokine over the number of tested pairs. Ribosomal proteins, defined using the CORUM database, showed significant enrichment as negative correlations passing ≤10% FDR. To test reproducibility across donors, the GDF6-ribosomal protein correlations were recomputed within each donor’s B cells separately using the exact Spearman method, requiring ≥30 co-observations per correlation. The correlation vectors between GDF6 and all ribosomal proteins per donor were then correlated using Spearman. To check whether this association was driven by cell size, PCA was computed using eigen decomposition of protein-correlations in the space of only ribosomal proteins, using the leading eigenvector as the ribosomal state axis. Each cell’s PC1 score along this axis was computed as the eigenvector-weighted sum of standardized ribosomal protein abundances, and correlated to GDF6 abundance across cells, and separately to the summed total protein content per cell. To functionally interpret the GDF6 covariation structure across B cells, partners were ranked by their GDF6 correlation and tested by pre-ranked gene-set enrichment (fgsea package, multilevel, gene-set sizes 5–300) against Reactome pathways (MSigDB C2 CP:REACTOME) and separately against the CORUM complex sets. Enrichment was assessed at ≤10% FDR, and redundant nested gene sets were collapsed for display, showing normalized enrichment scores. To check cell-type-specificity of the cytokine-ribosome anticorrelation, for each cell type, the cytokines showing the strongest negative ribosomal covariation were selected (prioritizing the count of significant negative ribosomal partners, falling back to the summed negative correlation), and their correlations against the ordered 40S and 60S subunits were displayed as a heatmap, with subunit column order derived from hierarchical clustering of the B cell block and significant associations (≤10% FDR) annotated.

## Data availability

Project meta data, processed data, and files required for reproducing all analyses are accessible via Zenodo DOI: 10.5281/zenodo.22649483.

## Code availability

Software, data-analysis pipelines and other supporting documentation are available at scp.slavovlab.net/2026_Khoury_et _al. The code for reproducing all the analyses and figures presented is freely available at GitHub (github.com/SlavovLab/PBMC_covariation).

## Acknowledgements

We thank V. Ondruska for help with timsTOF optimizations, and H. Specht and members of the Slavov laboratory for discussions and suggestions.

## Author contributions

Study design, supervision and raising funding: N.S. Data acquisition: L.K., S.K., A.L., M.H., J.E.M., J.M., Data analyses: L.K., J.E.M., and N.S., Initial draft: L.K. and N.S. Writing: all authors approved the final manuscript.

## Supplemental Figures

**Figure S1.**
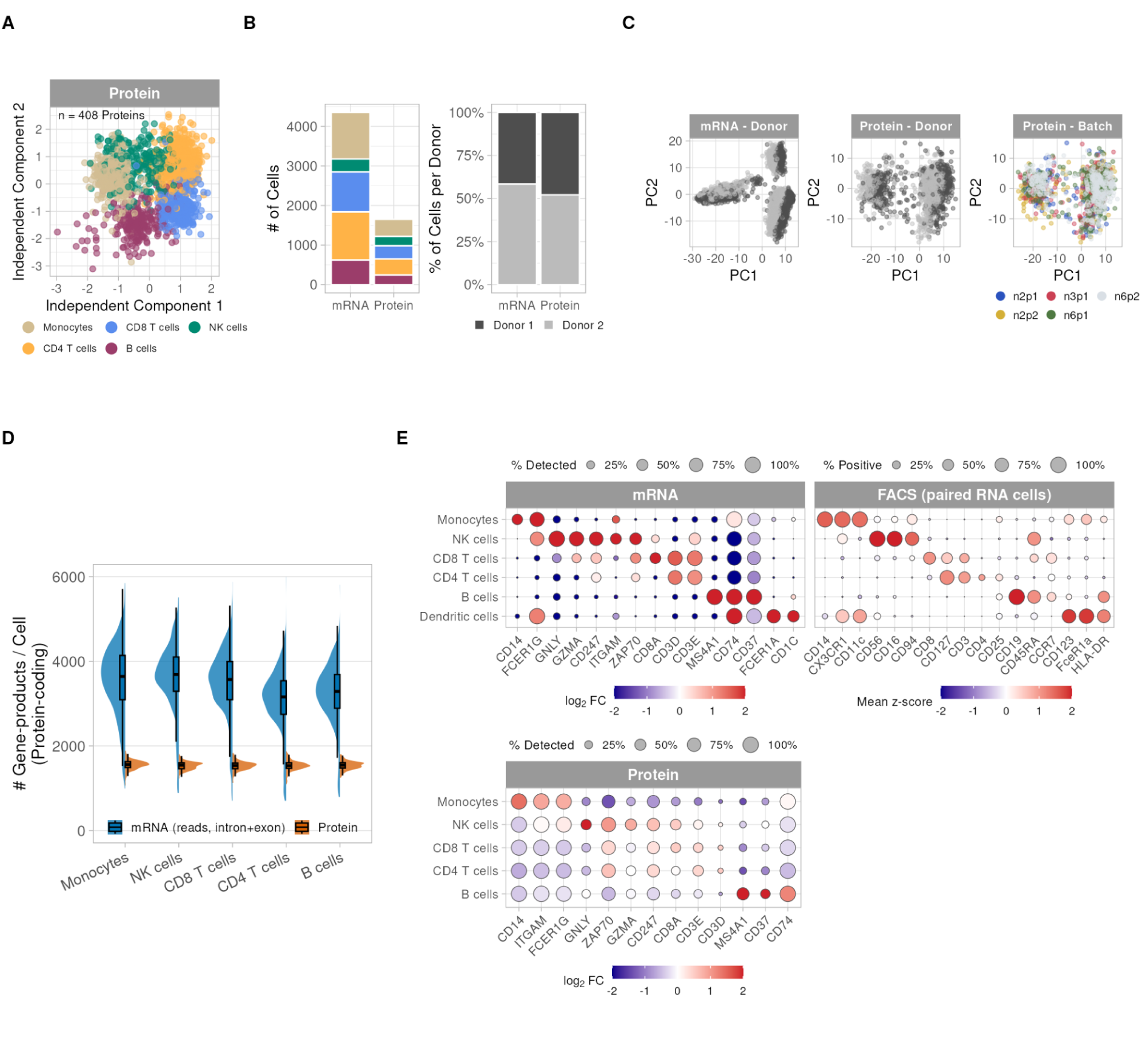
Cell type annotation across datasets and assessment of batch effects. **A** Independent component analysis of the protein dataset using only proteins quantified across all single cells (408 proteins), with cells colored by cell type. **B** Number of cells per cell type and proportion of cells per donor, by measurement modality. **C** Batch effects evaluated across mRNA and protein datasets after correction, visualized by PCA. **D** Quantified gene-products per cell across cell types, by modality, restricted to exon- and intron-containing protein-coding transcripts detected from read counts for the mRNA dataset. **E** Differential expression testing of marker gene-products across annotated cell types. Top: full mRNA dataset annotated using transcript abundances and for the subset of cells with paired FACS-based marker protein measurements from a fluorescent antibody panel (80.4% of cells). Bottom: protein dataset, using a highly overlapping marker set, based on differential protein abundance alone.

**Figure S2.**
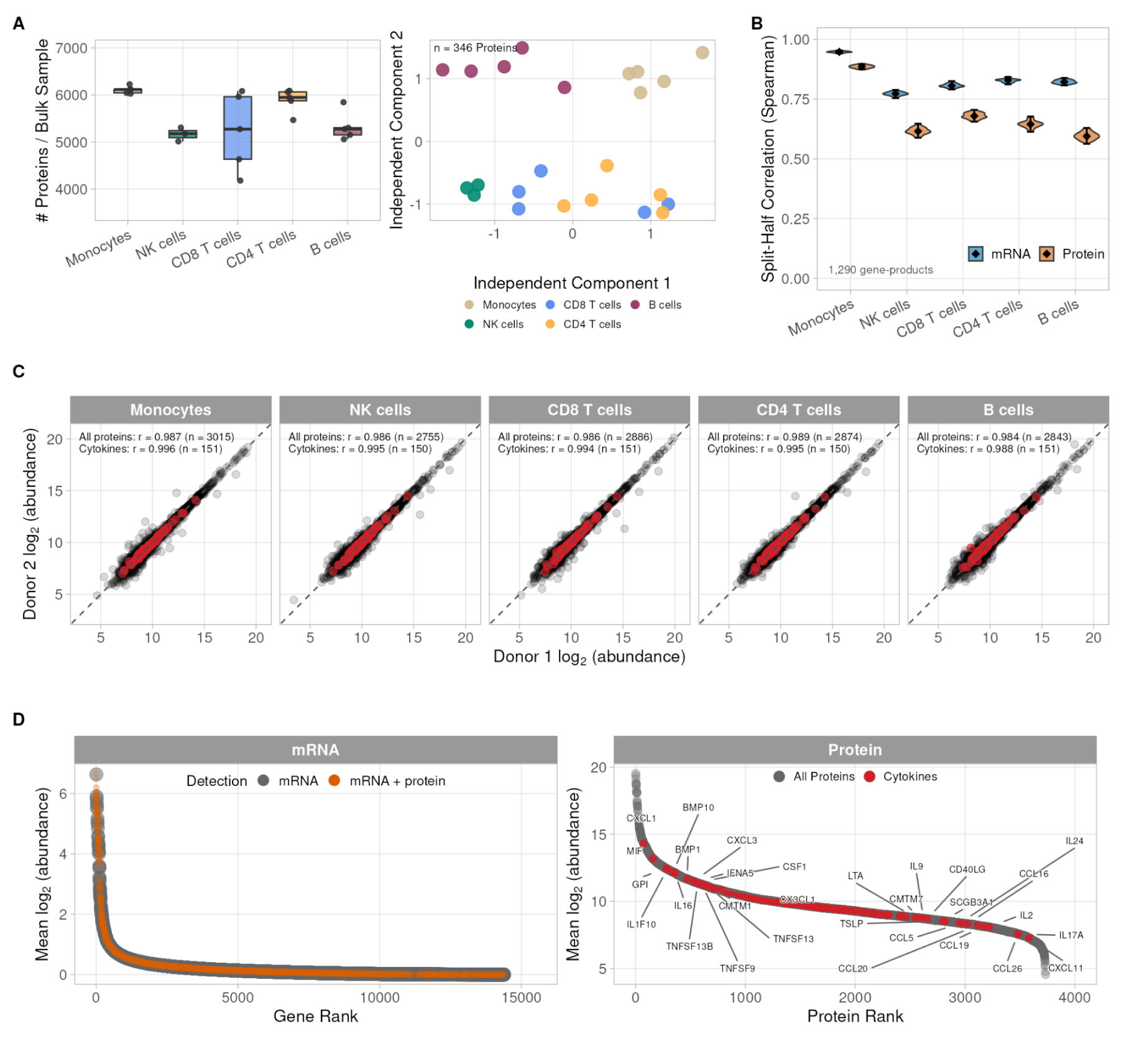
Dataset validation and depth comparison of the transcriptome and proteome across PBMCs. **A** Distributions of proteins quantified per bulk proteomic sample of FACS-isolated populations from the donors in this study by cell type (left) and ICA of bulk samples colored by cell type. **B** Split-half reproducibility of cell-type fold changes, in the feature space shared across modalities and cell types (1,290 gene-products) using downsampled numbers of cells from the mRNA data to match population sizes at the protein-level. For each cell type, cells of that type and the pooled remaining cells were each randomly split in half; the relative log2 abundance fold change of the cell type to all other cells was computed within each matched half, and the two fold change estimates were correlated across gene-products. Datapoints are mean Spearman correlations across 100 random splits; error bars are 95% confidence intervals. **C** Pearson correlation of log_2_ absolute protein abundances between donors within each cell type, for ∼2,700-3,000 proteins per cell type; the ∼150 quantified cytokines are shown separately. **D** Ranked mean transcript abundances from the Smart-seq3xpress dataset across quality-controlled cells and features, prior to filtering at <3% detection and <0.05 mean log-normalized abundance (n = 14,387 genes). Gene-products also quantified in the protein dataset (3,732 proteins) are highlighted in orange. **E** Ranked mean protein abundances across single cells, with quantified cytokines highlighted and the 15 most and least abundant cytokines annotated.

**Figure S3.**
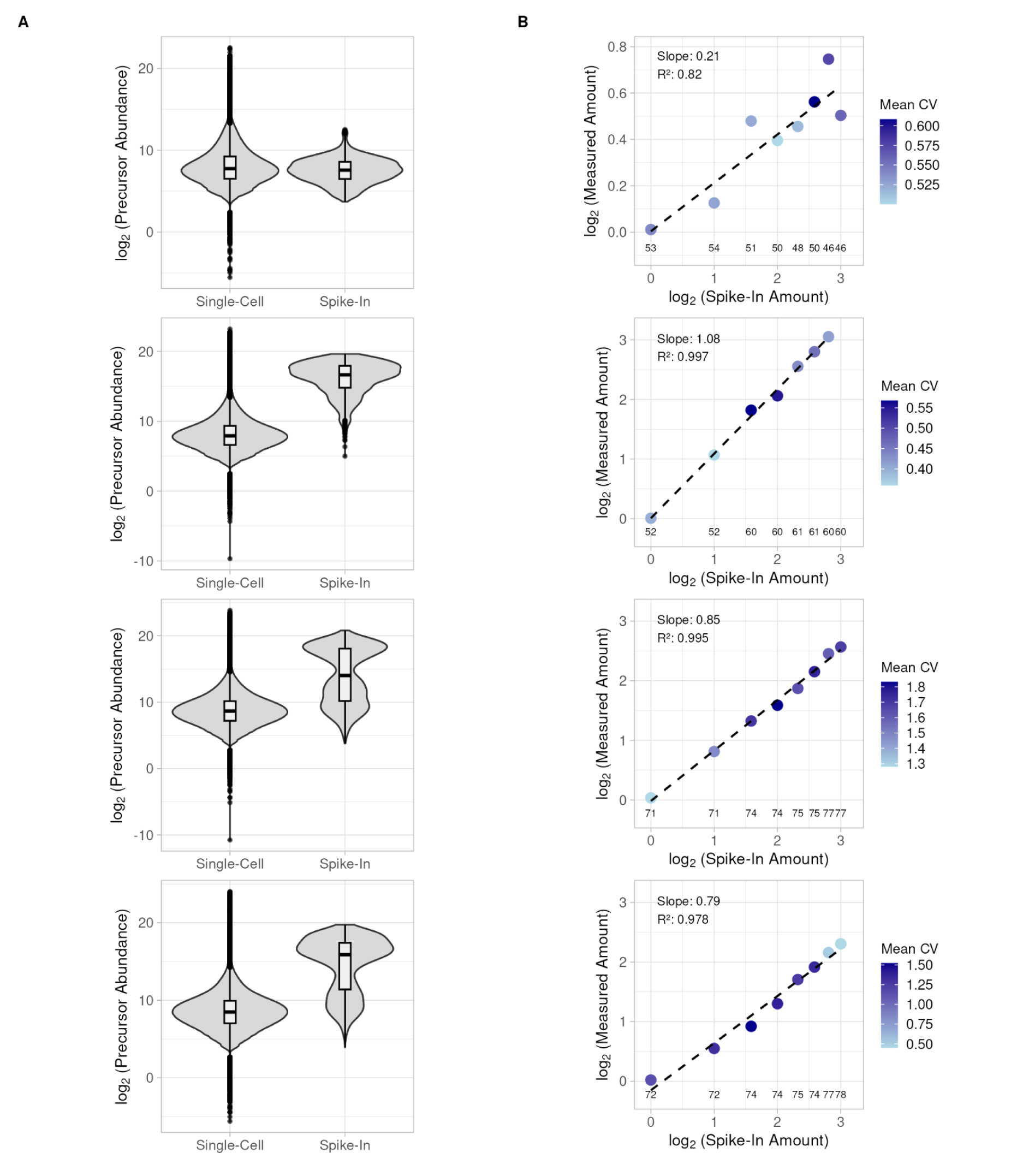
Quantitative accuracy of protein measurements assessed with a spike-in ladder of synthetic yeast peptides during the nPOP protocol. **A** Abundance distributions of precursor ions from endogenous PBMC peptides and from spike-in synthetic peptides, per nPOP prep. Spike-in amounts were designed to span the endogenous single-cell abundance range. Dataset 1 (top row) versus Dataset 2-4 (bottom three rows) differ in nPOP protocol version (REF). **B** Measured spike-in abundances, normalized to the lowest ladder level, versus known input amounts, per dataset. Lines are robust regression fits (rlm); R² calculated as the squared Pearson correlation between known and measured abundances.

**Figure S4.**
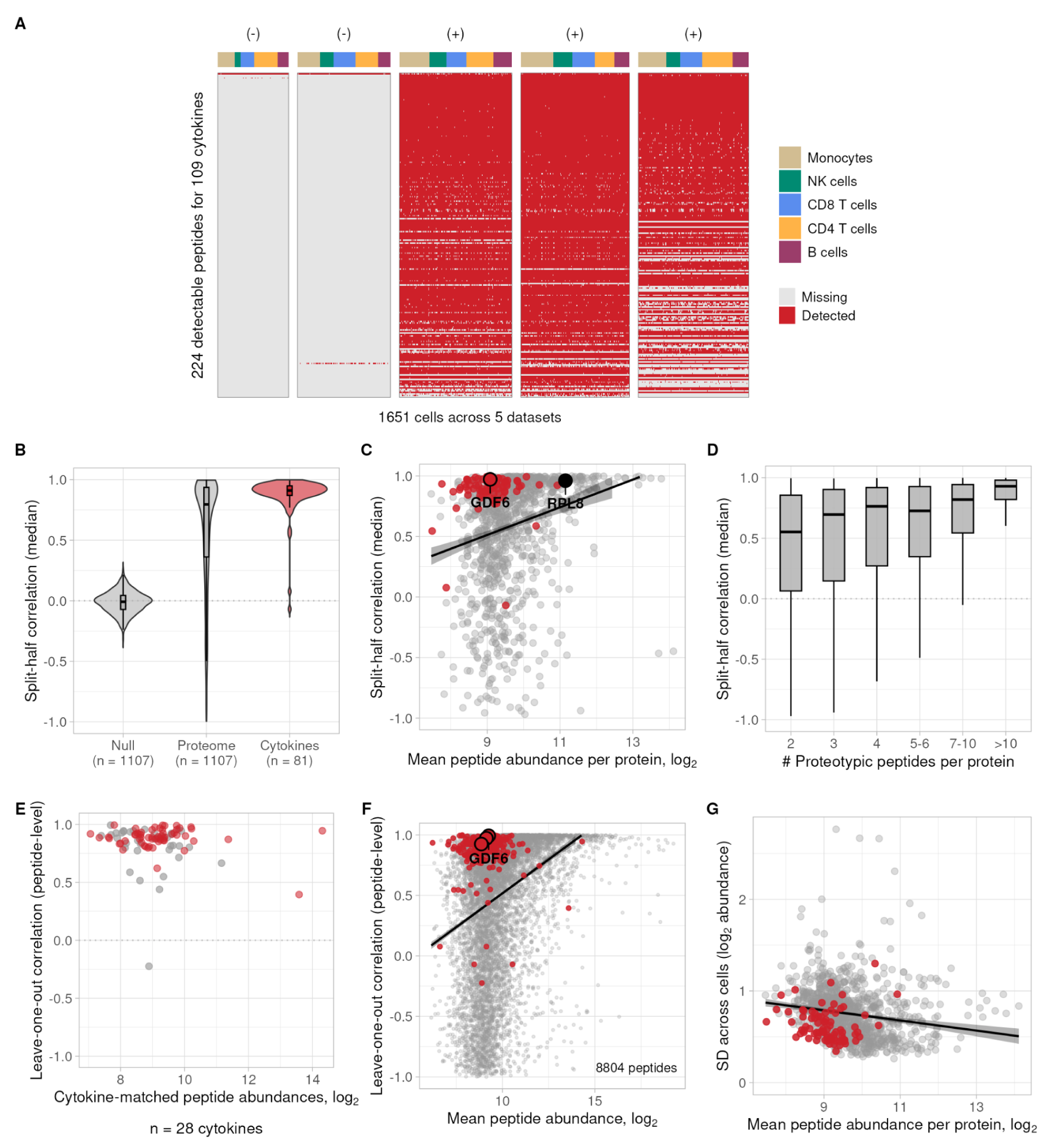
Protein quantification reliability across the proteome and for cytokines in the low-abundance regime in PBMCs. **A** Detection frequency of cytokine-mapping peptides included in the isotopologous carrier panel, in the three datasets where the panel was included versus the two where it was not. **B** Distributions of protein reliability estimates (median split-half correlation of proteotypic peptide sets, ≥2 peptides per protein; see Methods) for the full proteome and for quantified cytokines, with an empirical null from permuted cell-type labels. **C** Protein reliability versus mean peptide abundance per protein. GDF6 and RPL8 are labeled occupying similar reliability but different abundance regimes. **D** Protein reliability versus number of proteotypic peptides per protein. **E** Peptide-level reliability, estimated as leave-one-out correlation against the mean profile of remaining peptides mapping to the same protein (see Methods), for cytokine peptides included versus not included in the carrier panel, shown against peptide abundance. **F** Peptide reliability versus mean peptide abundance; three peptides mapping to GDF6 are labeled. **G** Standard deviation of log_2_ absolute protein abundances versus mean peptide abundance per protein. Line is a linear regression fit; cytokines are highlighted occupying the low abundance and low variance regime.

**Figure S5.**
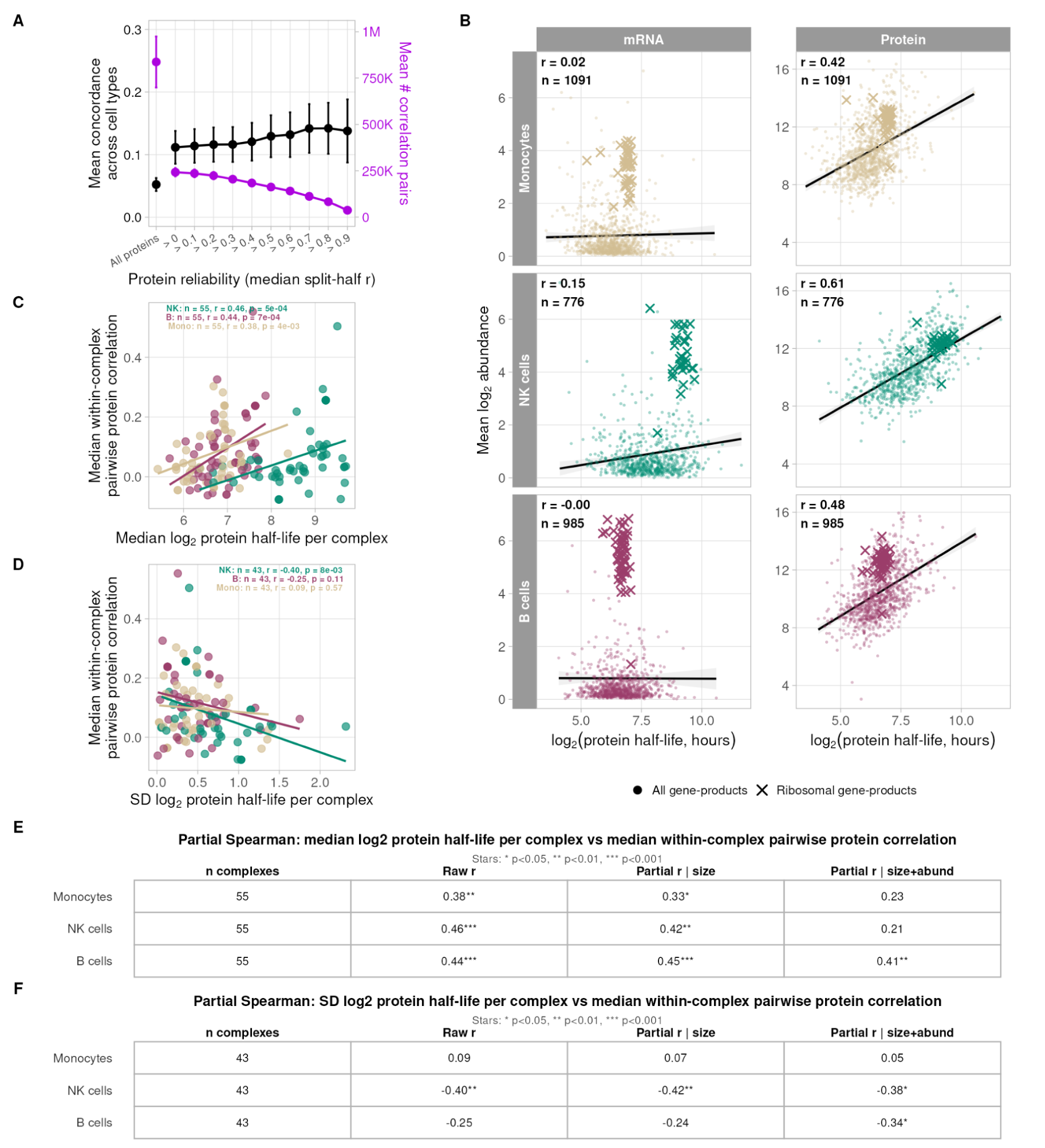
Concordance between mRNA and protein covariation networks and association of protein half-life with complex coordination across cell types. **A** Concordance between mRNA–mRNA and protein–protein correlations for shared gene-product pairs, per cell type, as a function of minimum protein reliability threshold (0.1 increments). Points are mean concordances and number of gene-product pairs per threshold in per cell type with standard deviation shown. **B** Correlation between protein turnover estimates (REF) within matched cell types at the protein- and mRNA-level. **C** Median within-complex pairwise protein correlation versus median log_2_ subunit half-life, per cell type, for complexes quantified across all cell types (n = 55 complexes). **D** Median within-complex pairwise protein correlation versus standard deviation of log_2_ subunit half-lives, per cell type, same complex set. NK cells show the largest effect size. **E** Partial correlations between median within-complex pairwise protein correlation and median log_2_ subunit half-life, conditioned on complex size (number of subunits) and on complex size plus median log_2_ subunit abundance, per cell type. **F** As in E, for the standard deviation of log_2_ subunit half-lives.

**Figure S6.**
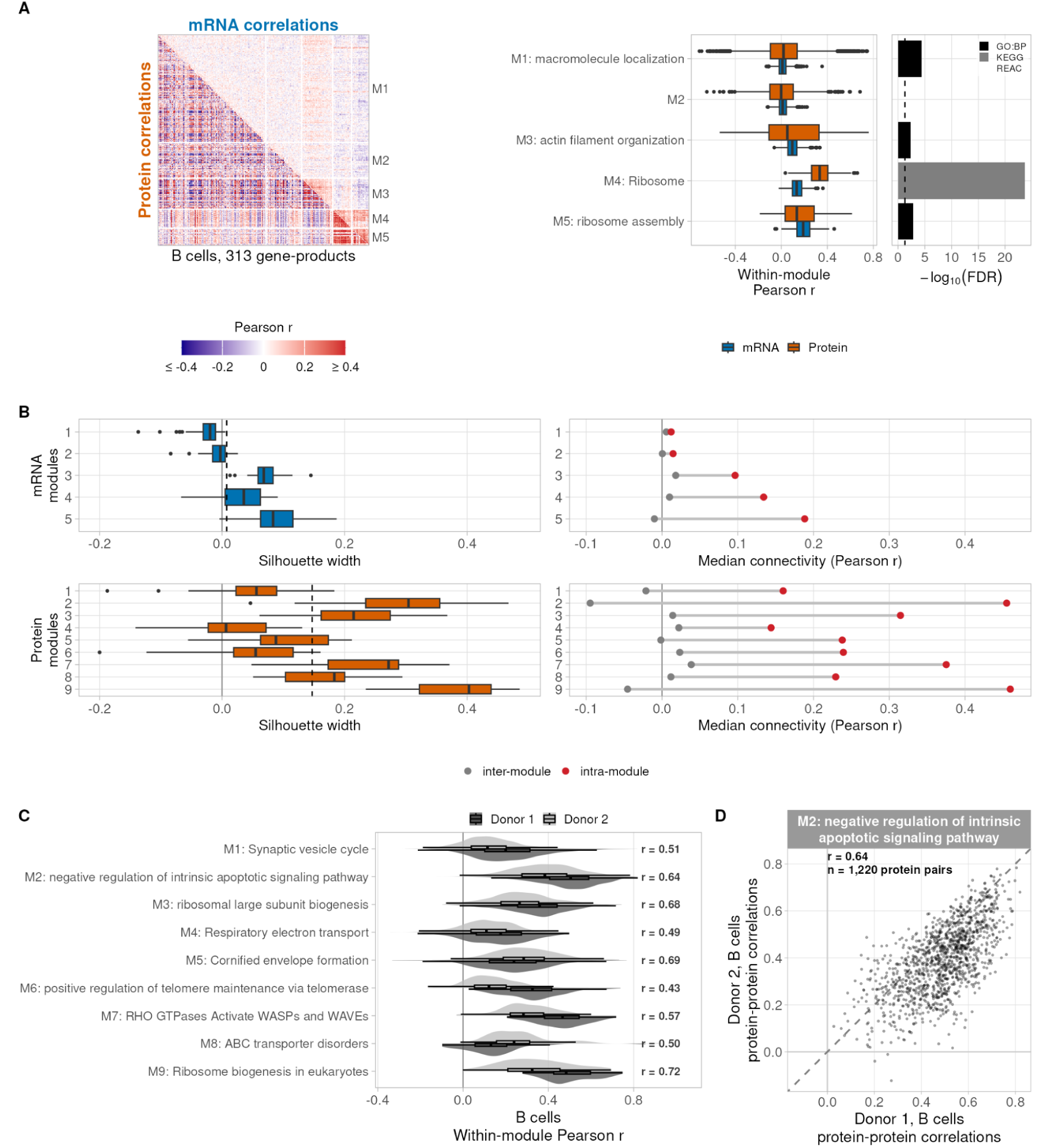
mRNA correlation modules within B cells and validation of correlation modules. **A** Correlation modules within B cells for the unified gene-product set, clustered hierarchically on mRNA correlation structure and annotated by enrichment analysis (GO, Reactome, KEGG; dashed line of 5% FDR shown), as in Fig. 2H. **B** Silhouette width per module and intra- versus inter-module connectivity of pairwise correlations, for protein and mRNA modules under their respective clustering conditions in the unified gene-product space. **C** Distributions of within-module protein correlations computed separately for each donor, using modules defined in Fig. 2H. Pearson correlations are annotated for each module across the pairwise complete set of protein-protein correlations between donors. **D** Scatterplot and concordance of Module 2 protein-protein correlations between donors computed independently.

**Figure S7.**
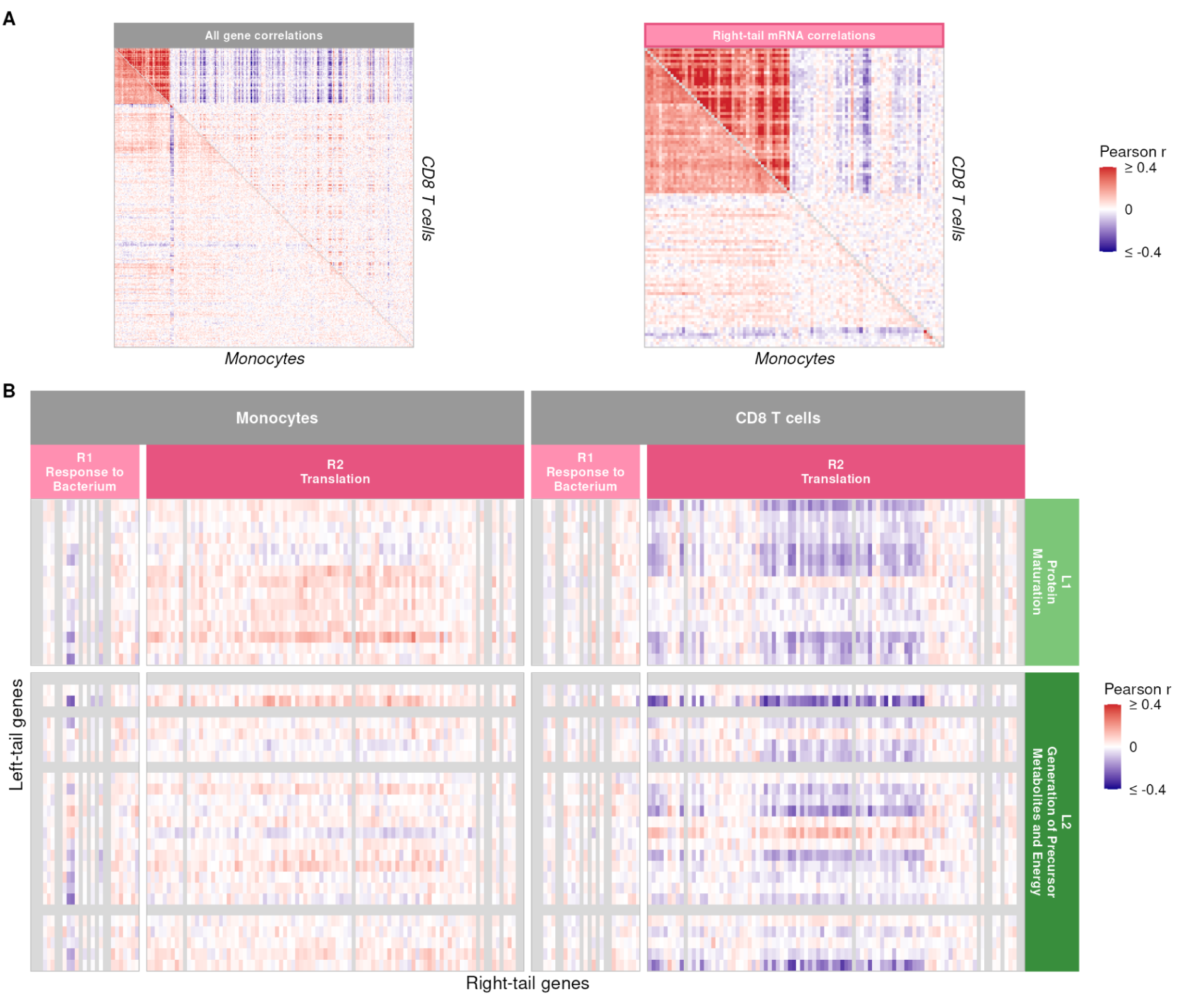
mRNA correlation structure for gene-products shared with the protein covariation analysis, monocytes and CD8 T cells. **A** mRNA–mRNA correlations for the gene-products intersected from the protein correlation vector analysis ordered by independent clustering. **B** Correlation heatmap of module blocks at the mRNA-level equivalent of Fig. 3B.

